# SARS-CoV-2 5’-UTR stem-loops activate the antiviral protein oligoadenylate synthetase 1 (OAS1)

**DOI:** 10.64898/2026.04.01.715984

**Authors:** Alejandro Oviedo, Camden R. Bair, Alexander P. Vasilakopoulos, Kierra Regis, David Vanlnsberghe, Anice C. Lowen, Graeme L. Conn

## Abstract

The innate immune system relies on pathogen recognition receptors, such as the 2’,5’-oligoadenylate synthetase (OAS) proteins, to detect pathogen-associated molecular patterns like viral double-stranded (ds)RNA. A specific splicing variant of OAS1 (OAS1-p46) has been implicated in initiating an immune response that leads to decreased disease severity during SARS-CoV-2 infection. OAS1-p46 has a C-terminal lipid modification motif that allows for anchoring of the protein to intracellular membranes and thus potential colocalization with immunogenic viral RNA regions such as the SARS-CoV-2 5’-untranslated region (5’-UTR). Here, we show that OAS proteins can detect the 5’-structured elements (5’-SE)–comprising the 5’-UTR and three additional stem-loop structures–and activate the cellular RNase L pathway. Through systematic 3’-end truncations of the 5’-SE, we show that the smallest 5’-SE fragment capable of potently activating OAS1 is the conserved and highly structured SL1-4b region, containing the first four stem-loop structures and their intervening linking sequences (SL1-4), as well as an unstructured region (“SL4b”) to the 3’-side of SL4. Analyses of OAS1 activation and RNA secondary structure probing using selective 2’-hydroxyl acylation analyzed by primer extension and mutational profiling (SHAPE-MaP) of additional SL1-4b RNA variants suggests a model in which SL4 acts as the primary OAS1 interaction site, while the unstructured SL4b region and two other stem-loops (SL1 and SL3) are necessary for optimal presentation of this region for OAS1 activation. Our findings reveal a structurally complex viral RNA region that potently activates OAS1, underscoring the potential complexity of RNAs that can strongly activate this innate immune sensor.

## Introduction

The innate immune system is a critical line of cellular defense that uses pattern recognition receptors (PRRs) to sense a variety of pathogen-associated molecular patterns (PAMPs) (1-7). The oligoadenylate synthetase (OAS) family of nucleotidyl transferases comprises an important group of interferon-stimulated PRRs that detect dsRNA, a potent PAMP of viral infection (7-10). Upon binding dsRNA, the OAS proteins synthesize the 2’,5’-linked oligoadenylate second messenger which, when >3 adenylates in length, promotes dimerization and activation of the latent ribonuclease RNase L, and subsequent cleavage of viral and cellular RNA. Activation of the OAS/RNase L pathway thereby promotes a translational control program in the infected cell, limiting viral replication (9-13).

The OAS protein family comprises three catalytically active members, OAS1, OAS2, OAS3, and the catalytically inactive OASL, which also functions in innate immune sensing but acts independently of RNase L (14-20). OAS3 has the highest affinity for dsRNA and detects longer dsRNA regions (21-23), making it the primary family member responsible for detection of many viral pathogens (24,25). However, unlike OAS1 and OAS2, which have isoforms that are localized to membrane-bound organelles (25-28), OAS3 is thought to be predominantly cytosolic, rendering it unable to detect viral RNAs that are sequestered within such compartments (25,29). Human OAS1 has at least four isoforms stemming from alternative splicing of the *OAS1* gene (15,30-32). These OAS1 isoforms share a common core (amino acids 1-346) with splicing affecting only the C-terminal sequence and altering localization without impacting catalytic activity (30,33-35). The two major isoforms are OAS1-p42, which is predominantly cytosolic, and OAS1-p46, which is C-terminally lipid-modified and membrane anchored. These two isoforms arise due to a single nucleotide polymorphism (SNP) in *OAS1*, which shifts the splice-acceptor site in OAS1 exon 6 (25,32). Individuals with at least one G allele (SNP 12-112919388-G-A) produce OAS1-p46 which contains a canonical CAAX (cysteine/ aliphatic/ aliphatic/ any)-box prenylation signal that allows anchoring of the lipid-modified OAS1 into the cytosolic face of intracellular membranes (30,36,37). As a result, OAS1-p46 shows preferential antiviral activity against viruses that use membrane-bound replication organelles, such as encephalomyocarditis virus, West Nile virus and SARS-CoV-2 (25,32,38,39).

SARS-CoV-2 is a positive-sense RNA virus and the causative agent of COVID-19 disease (40,41). COVID-19 disease severity and patient outcome was found to be dependent on the same *OAS1* SNP that results in production of OAS1-p46, i.e. individuals with OAS1-p46 had more favorable outcomes compared to those who express only OAS1-p42 (38,42). This finding led to the proposal that, during SARS-CoV-2 replication, dsRNA intermediates are accessible to the membrane-anchored OAS1-p46, but not cytosolic OAS proteins, resulting in activation of the OAS/RNase L pathway (38). As part of its ∼30 kb RNA genome, the first 480 nucleotides–encompassing a 5’-untranslated region (5’-UTR) and part of the first open reading frame (ORF1a)–form a highly structured domain known as the 5’-structural elements (5’-SE; **Fig.1**). The 5’-SE comprises eight stem-loop (SL) structures with SL1-4 and part of SL5 forming the 5’-UTR and the remainder of SL5 and SL6-8 encoding the start of ORF1a. The coronaviral 5’-UTR is functionally important for regulation and efficiency of viral RNA synthesis (39,43-49). Individual nucleotide-resolution cross-linking and immunoprecipitation (iCLIP2) applied to infected cells with exogenously expressed OAS proteins revealed that SL1 and SL2 most prominently immunoprecipitated with OAS1 (38). SL1 and SL2 were thus proposed as individual binding sites for OAS1, prompting another study modeling a putative OAS1-SL1 complex (50). However, this model was not experimentally tested and the base-paired regions of both SL1 and SL2 are substantially shorter than the minimum ∼17 base pairs (bp) needed for OAS1 activation (23,35). As a result, the region of the SARS-CoV-2 5’-SE responsible for activation of OAS1 remains undefined.

**Fig. 1.**
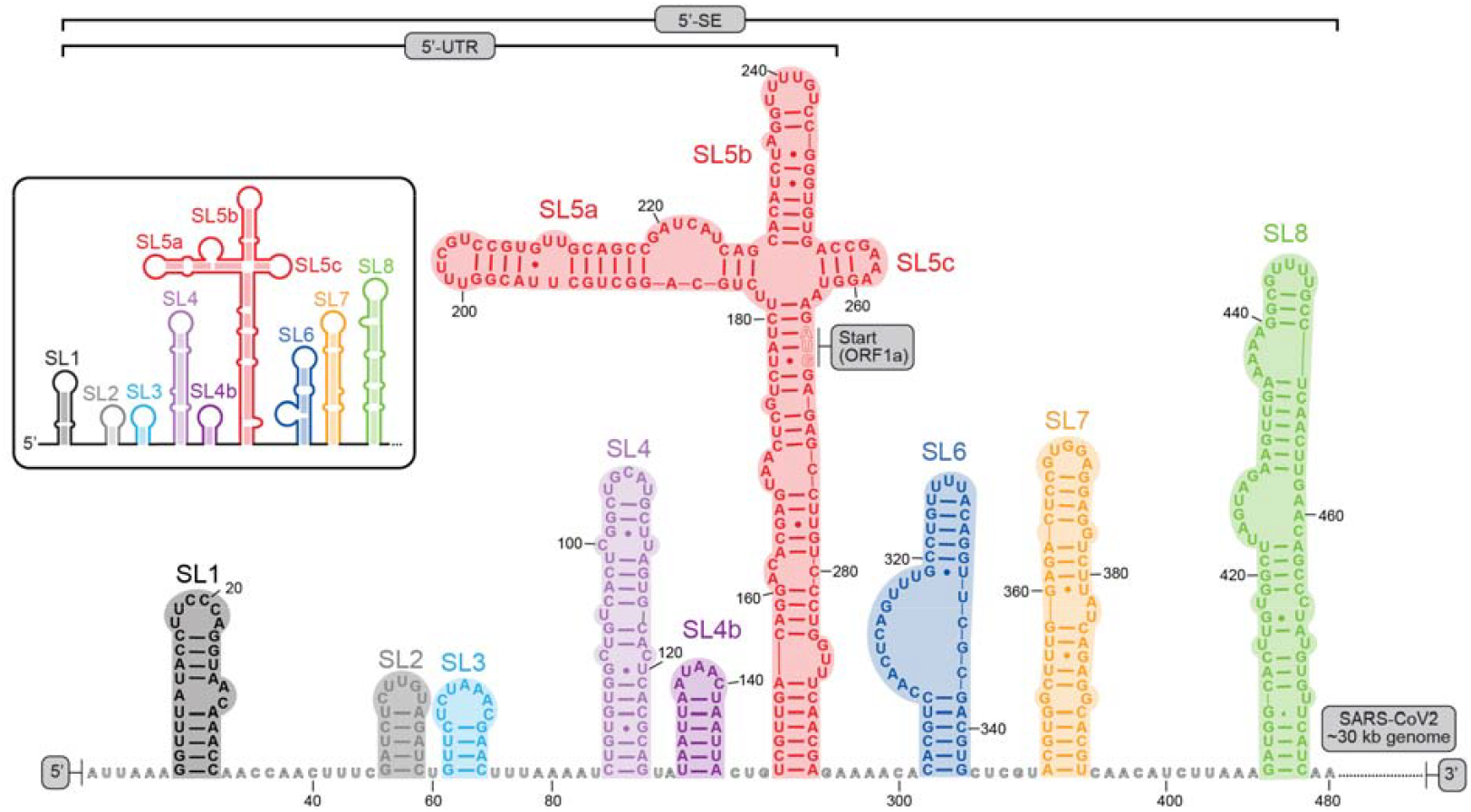
Sequence and secondary structure of the SARS-CoV-2 genomic RNA 5’-end. Sequence and secondary structure of the overlapping SARS-CoV-2 5’-UTR and 5’-SE regions. Individual stem-loops are color-coded and the full structure shown as a cartoon representation (*inset*) as used in later figures.

Here, we show that although the full SARS-CoV-2 5’-SE (SL1-8) RNA potently activates OAS1, neither SL1 nor the combination of SL1-2 is sufficient for this activity. Through systematic truncation of the 5’-SE, we identify a larger region comprising SL1 to SL4b (SL1-4b) as the minimal fragment needed for full OAS1 activation. Through further dissection of the necessary components of SL1-4b for OAS1 activation, we identify the primary importance of SL4 but also find that multiple features of the SL1-4b region contribute to the ability of this RNA region to activate OAS1. This work thus broadens our understanding of how dsRNA is sensed by OAS1 in the context of structured domains and underscores the potential complexity of RNAs that can strongly activate this innate immune sensor.

## Results

### The SARS-CoV-2 5’-SE RNA strongly activates the OAS family proteins

To test whether the SARS-CoV-2 genomic RNA 5’-SE activates the cellular OAS/RNase L pathway, we cloned the full region (SL1-8, nucleotides 1-480; **Fig. 1**) into a plasmid designed for *in vitro* transcription (IVT) (51). RNAs were synthesized with an appended hepatitis delta virus (HDV) ribozyme to ensure homogenous 3’-ends and to enable resolution during purification from potential immunogenic dsRNA contaminants. The purified SL1-8 RNA (**Supplemental Fig. S1A**) was transfected into wild-type, OAS3 knock-out (KO), and RNase L KO A549 cells pre-treated with interferon (IFN)-β1α. OAS/RNase L pathway activation was then assessed via Bioanalyzer analysis of 28S and 18S rRNA integrity (**Fig. 2A-C**) in comparison to mock transfection (negative control) and transfection with the dsRNA analog polyinosinic-polycytidylic acid (poly(I:C); positive control). For each RNA, band intensities were also quantified for cleaved and uncleaved 28S rRNA from three independent experiments, and expression or absence (in KO cells) of OAS3 or RNase L was confirmed by immunoblotting (**Fig. 2D**). Transfection of SL1-8 RNA resulted in clear cleavage of 28S and 18S rRNA in wild-type cells. In OAS3 KO cells, cleavage of 28S and 18S was also observed, albeit at a reduced level (with ∼50% lower cleaved product similar to the reduction observed for poly(I:C)). In contrast, minimal cleavage products were observed in RNase L KO cells, confirming that rRNA degradation in wild-type and OAS3 KO cells was due to OAS/RNase L pathway activation.

**Fig. 2.**
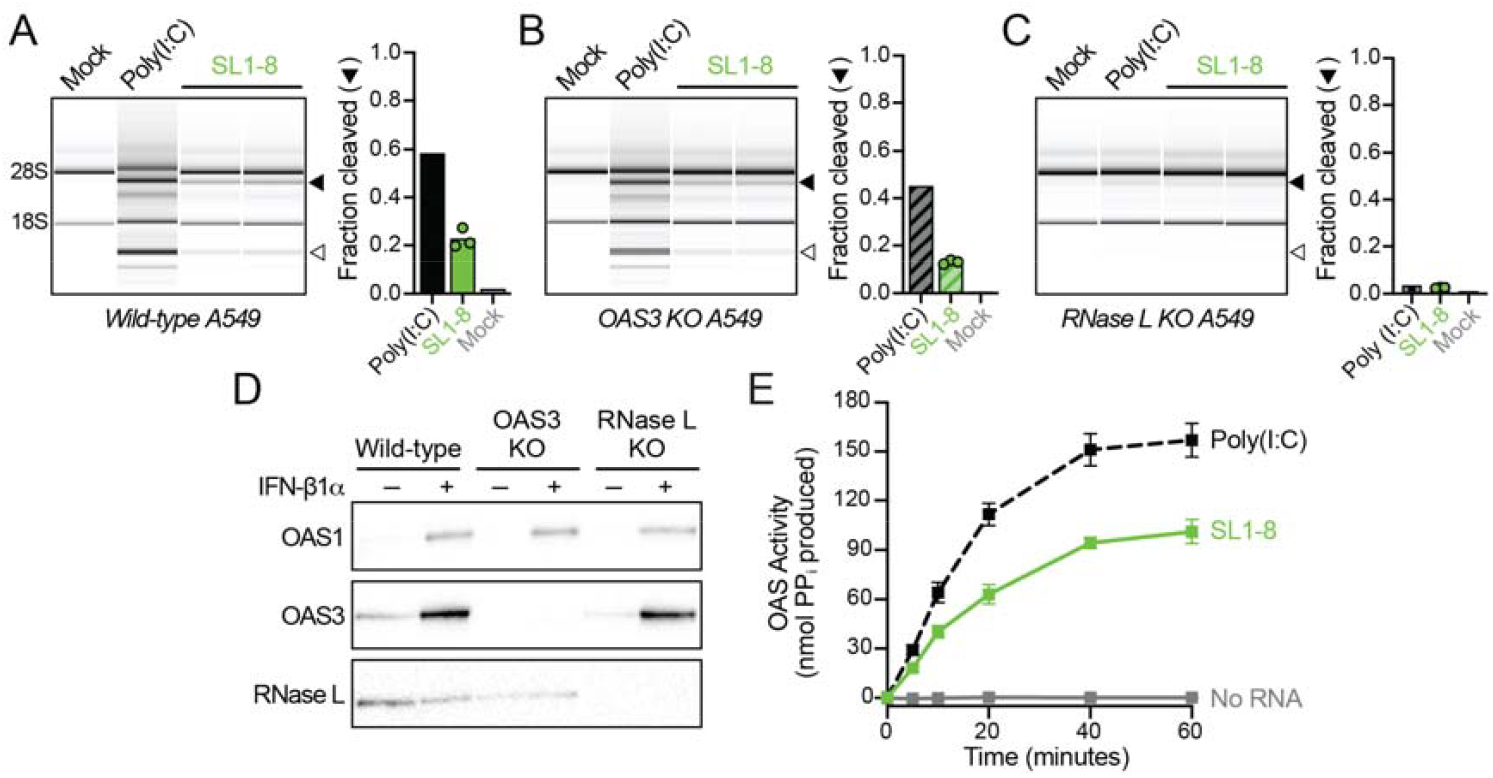
SL1-8 of the SARS-CoV-2 5’-SE activates OAS proteins in cells and *in vitro*. Bioanalyzer analyses of rRNA integrity in **(A)** wild-type, **(B)** OAS3 KO, and **(C)** RNase L KO A549 cells following transfection with SL1-8 RNA. Cleavage products of 28S and 18S are indicated by the solid and open arrows, respectively. Mock and poly (I:C) serve as negative and positive controls. Band intensities for cleaved and uncleaved 28S were quantified for three independent transfections and were used to calculate fraction cleaved. The extent of 28S rRNA cleavage was quantified for three independent transfections of each cell type as the ratio of band intensities for the cleaved (solid arrow) to uncleaved bands. **(D)** Immunoblots confirming expression of OAS1 (top) after IFN-β1α treatment in wild-type, OAS3 KO, and RNase L KO cells. OAS3 KO and RNase L KO are confirmed in middle and bottom blots, respectively. **(E)** *In vitro* chromogenic OAS1 activity demonstrating that SL1-8 potently activates OAS1, with poly(I:C) and no RNA reactions serving as positive and negative controls, respectively.

These results thus reveal OAS3 to be strongly activated by SARS-CoV-2 SL1-8 RNA when transfected into the cytoplasm, where OAS3 is exclusively expressed. Further, these results confirm that OAS1 and/or OAS2 are also activated by SL1-8 in a cellular context. To confirm specific activation of OAS1 by SL1-8 RNA, we turned to an established *in vitro* activity assay (34,52,53) using purified recombinant OAS1 (“core” residues 1-346; **Supplemental Fig. S1B**) since the C-terminal sequence only affects localization and not activity of the protein. These analyses confirmed that OAS1 is potently activated by SL1-8 RNA (**Fig. 2E**).

### An extended region of the SARS-CoV-2 5’-SE is required for full OAS1 activation

Previous studies implicated SL1 and SL2 as individual activating elements within the 5’-UTR, but did not directly test the capacity of these RNA regions to activate OAS1 (38,50). Importantly, the dsRNA helical regions of SL1 and SL2 are ∼11 bp (including bulge nucleotides) and 5 bp, respectively (**Fig. 3A**), and are thus substantially shorter than the established ∼17 bp minimum dsRNA length requirement for OAS1 activation (35). Consistent with this, we used a chemically synthesized SL1 RNA construct in the OAS1 *in vitro* activity assay and observed no activation (**Fig. 3B**). We speculated that SL1 and SL2 could potentially stack coaxially to make a ∼17 bp region, meeting the required dsRNA length for OAS1 activation (**Fig. 3A**, *inset*). We therefore generated two RNA IVT constructs: one containing the full 5’-end single-stranded sequence (SL1-2) but with the 5’-end modified to incorporate two G nucleotides for transcription by T7 RNA polymerase (RNAP), and a second (SL1-2 (Δ5’)) with this 5’-single-stranded region deleted to exploit the GG dinucleotide at the base of SL1 for transcription (**Fig. 3A**). Again, both SL1-2 and SL1-2 (Δ5’) RNAs failed to activate OAS1 in the *in vitro* activity assay (**Fig. 3B**).

**Fig. 3.**
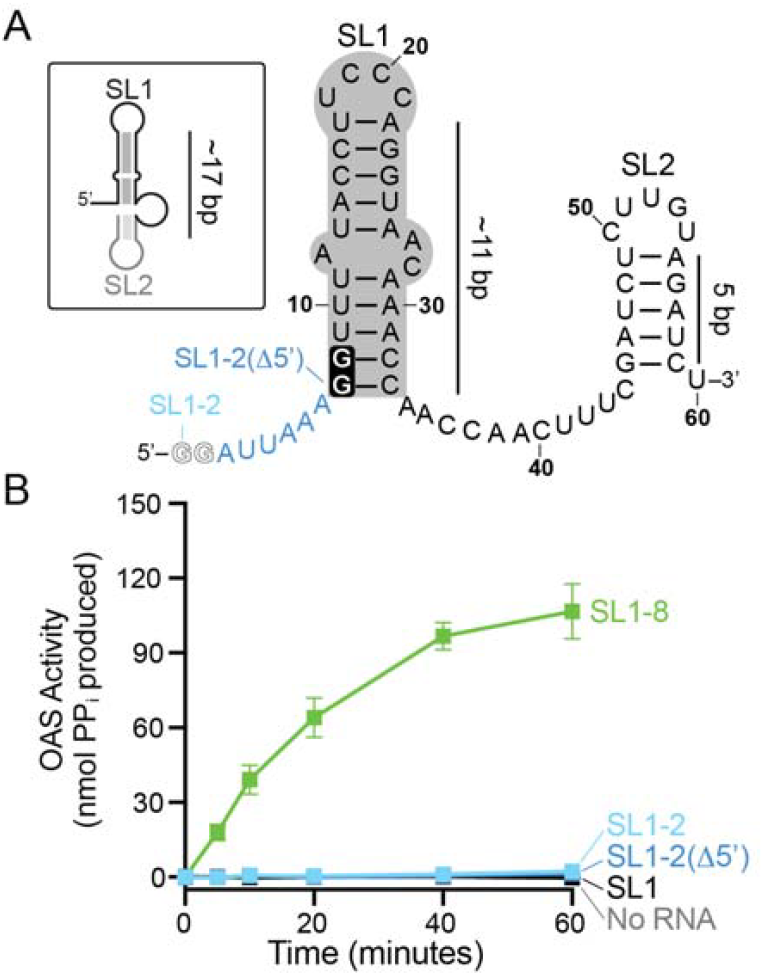
Isolated SL1 and SL1-2 RNA regions from the SARS-CoV-2 5’-SE do not activate OAS1. **(A)** Sequence and secondary structure of SL1 (gray shading) and SL2 of the SARS-CoV-2 5’-SE and their approximate double-stranded helical lengths in base pairs (bp). Also indicated are the 5’-GG dinucleotides added to the 5’-end single stranded region (light blue; GG in outline font) and at the base of SL1 (black background) for IVT of SL1-2 and SL1-2(Δ5’), respectively. Boxed (*top left*), cartoon depicting potential SL1/SL2 coaxially stacking to form a continuous dsRNA helix of sufficient length to bind and activate OAS1 (∼17 bp). **(B)** *In vitro* chromogenic assay showing that OAS1 is not activated by SL1, SL1-2, or the SL1-2 region lacking the single-stranded 5’-end sequence (SL1-2(Δ5’)).

Since SL1-2 alone did not activate OAS1, we next sought to identify the minimal region within SL1-8 required for OAS1 activation by making new constructs via systematic SL truncations from the 3’ end (**Fig. 4A** and **Supplemental Fig. S1A**) and testing the ability of each RNA to activate OAS1 in the *in vitro* activity assay (**Fig. 4B**). These analyses revealed SL1-4b to be the shortest 5’-SE fragment that activates OAS1 to the same extent as SL1-8. SL1-4 exhibited an intermediate capacity to activate OAS1, while all activation was eliminated with further truncation (**Fig. 4B**). Given that SL4 is >17 bp and thus of sufficient length for OAS1 activation, these results indicate that SL4 likely serves as the main interaction site for OAS1, with SL4b further promoting OAS1 activation. While SL4b is too short to individually promote OAS1 activation (and may not form a stable helical structure–see below), precedence for additional sequence or structural features enhancing OAS1 activation by a dsRNA region comes from our prior studies of the 3’-single stranded pyrimidine (3’-ssPy) motif and the cellular non-coding RNA nc886 (53,54).

**Fig. 4.**
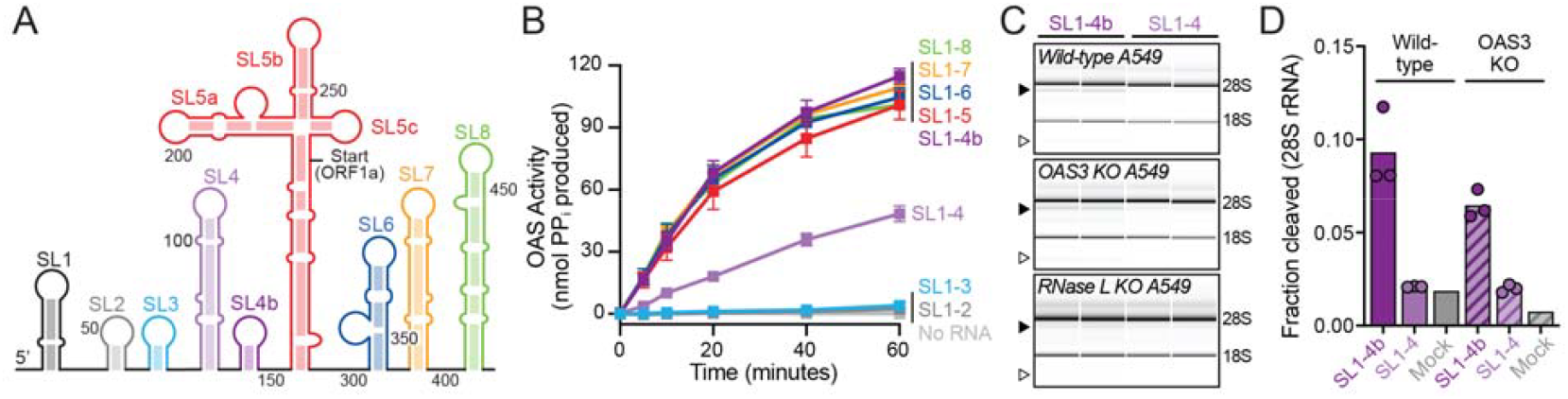
Systematic truncation of the 5’-SE reveals SL1-4b as the shortest section that potently activates OAS1. **(A)** Cartoon of first 8 stem-loops (5’-SE) of the SARS-CoV-2 genomic RNA. **(B)** *In vitro* OAS1 activity assay showing that truncation of 3’-end stem-loops does not decrease activation of OAS1 until SL4b is removed, making SL1-4b the shortest section that fully activates OAS1. SL1-4 exhibits intermediate capacity to activate OAS1, while SL1-3 and SL1-2 fail to activate. **(C)** Bioanalyzer analysis of 28S and 18S integrity in wild-type, OAS3 KO, and RNase L KO A549 cells following transfection with SL1-4b or SL1-4. Cleavage products of 28S and 18S are shown by the black and white arrows, respectively. SL1-4b transfection results in more 28S cleavage product than SL1-4 mirroring *in vitro* OAS1 activation. **(D)** Fraction of cleaved 28S was quantified for three independent transfections of each RNA in wild-type and OAS3 KO cells, confirming that SL1-4b more potently activates the OAS/RNase L pathway than SL1-4.

Next, we asked whether these shorter 5’-SE fragments could activate the OAS/ RNase L pathway in the cell, as observed above for SL1-8. SL1-4 and SL1-4b RNAs were transfected into wild-type, OAS3 KO, and RNase L KO A549 cells pre-treated with IFN-β1α, as before, and pathway activation was assessed via Bioanalyzer analysis of 28S and 18S rRNA cleavage. Consistent with their *in vitro* activities, SL1-4b exhibited stronger OAS/RNase L pathway activation, as indicated by greater 28S rRNA cleavage, compared to SL1-4 in both wild-type and OAS3 KO cells (**Fig. 4C,D**). Minimal cleavage was detected in RNase L KO cells, as anticipated. Although 28S rRNA cleavage is reduced in OAS3 KO cells for SL1-4b, this reduction is less pronounced than the reduction observed for SL1-8, suggesting that SL1-4b is more specific for the smaller OAS proteins than OAS3. These results validate SL1-4b as the shortest region capable of full OAS1 activation and demonstrate that when transfected into the cytoplasm, the SL1-4b RNA can potently activate OAS1/ OAS2 in the cellular context mirroring activation of OAS1 *in vitro*.

### SL4b does not form a stable SL but may alter the SL1-4 RNA structure in a manner necessary for full OAS1 activation

The SARS-CoV-2 5’-UTR structure has been examined in several studies using chemical probing approaches both in cell and *in vitro* (46,47,55-59). Interestingly, the resulting secondary structures for SL4 and SL4b differ in these studies with a SL structure proposed for SL4b in two cases (47,58), while others suggested that the region forms an unstructured expansion of the linker sequence between SL4 and SL5 (46,56,57,59) or an extension of SL4 (55). Given the importance of SL4b for OAS1 activation identified above, we used SHAPE-MaP to assess the SL4b structure within the SL1-4b fragment and to compare the structures of the other SLs in its presence and absence (i.e., between SL1-4b and SL1-4). Three independent modification experiments were performed using the 2A3 SHAPE reagent, and averaged normalized reactivities were plotted and mapped onto the predicted RNA secondary structures (**Fig. 5A,B**). The nucleotide reactivities verify the stable formation of each SL in the SL1-4 region in both RNAs with only minor differences in a small number of nucleotides. However, the uniformly high reactivity of the nucleotides proposed to form the 5’-half of SL4b clearly indicates that this region does not fold into a stable SL under the conditions used in our assay (**Fig. 5B**).

**Fig. 5.**
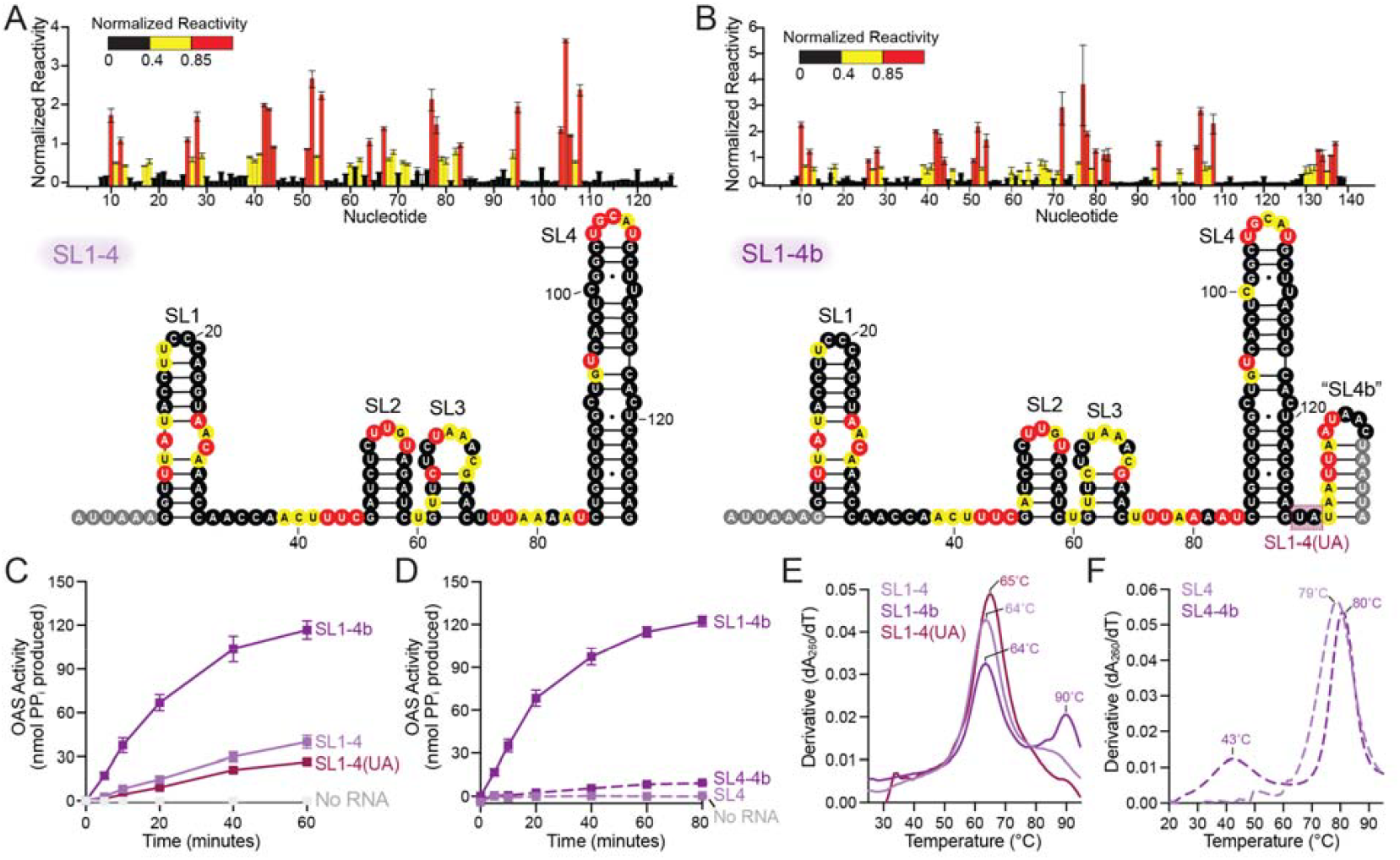
SL4b is unstructured but alters the structure and OAS1 activation capacity of the SL1-4 region. **(A)** Plot (*top*) and mapping onto the SL1-4 RNA secondary structure (*bottom*) of average normalized SHAPE reactivity for three independent 2A3 modification experiments. Nucleotide reactivity is consistent with the formation of each stem-loop structure. Gray shaded nucleotides indicate primer sites for which values could not be calculated. **(B)** As panel A but for SL1-4b RNA. Nucleotide reactivity is again consistent with the formation of SL structures in the SL1-4 region but does not support formation of a structured SL4b under the conditions used. The low reactivity of two nucleotides on the 3’-side of SL4 which prompted testing of an additional RNA construct, SL1-4(UA), is indicated (shaded red box). **(C)** *In vitro* OAS1 activity assay showing that SL1-4(UA) RNA promotes intermediate OAS1 activation, similar to SL1-4. **(D)** *In vitro* OAS1 activity assay showing that the isolated SL4 does not activate OAS1; addition of SL4b in this RNA context results in a very weak activator. **(E)** RNA UV thermal melting analysis of SL1-4b and SL1-4 in 100 mM NaCl, reveals an additional structural feature in SL1-4b that unfolds at higher temperature (T_m_ ∼95°C). **(F)** RNA UV thermal melting analysis of SL4 and SL4-4b in 100 mM NaCl and 7 mM MgCl_2_, revealing an additional low temperature apparent transition (T_m_ ∼45 ∨C) for SL4-4b that is not present for SL4, as well as a small increase (∼3 °C) in the T_m_ of the high temperature apparent transition.

In contrast to the high reactivity of SL4b, the dinucleotide (UA) linker between SL4 and SL4b exhibited low reactivity, suggesting its possible involvement in extending the SL4 structure or potentially enhancing activation by SL4 in some other manner, e.g. similar to the 3’-ssPy motif (53). To test this idea, we generated SL1-4(UA), ending the RNA with the dinucleotide sequence that follows SL4 (boxed region on the SL1-4b secondary structure in **Fig. 5B**). However, SL1-4(UA) promoted essentially the same intermediate level of OAS1 activation in the *in vitro* assay as SL1-4, indicating that additional nucleotides of the SL4b are required for full OAS1 activation despite its apparent lack of defined secondary structure (**Fig. 5C**).

Given that SL4 is >17 bp and thus sufficiently long to bind and activate OAS1, we next asked whether the isolated SL4-4b region is sufficient for OAS1 activation using the same *in vitro* assay. However, these analyses revealed the isolated SL4-4b RNA to be a very weak activator, while SL4 alone did not activate above the no RNA control under the standard assay conditions (**Fig. 5D**). This result indicates that other elements of the SL1-3 region contribute directly, or in some supporting role, to activating OAS1.

To gain further insight into whether the presence of SL4b impacts the structure or stability of the SL1-4 region, we used UV thermal melting analysis to assess RNA unfolding under three conditions related to those used in the *in vitro* OAS1 activation assay, as we have done previously for a short model dsRNA OAS1 activator (60): low salt (10 mM NaCl), high salt (100 mM NaCl), and high salt with Mg^2+^ (100 mM NaCl and 7 mM MgCl_2_) (**Fig. 5E,F** and **Supplemental Fig. S2A-C**). SL1-4 and SL1-4b unfold with similar profiles and a major apparent unfolding transition with T_m_ of ∼64 °C under the high salt condition (**Fig. 5D**). However, SL1-4b also displays a second apparent unfolding event at higher temperature (Tm ∼90 °C), which is absent in SL1-4, that may reflect an additional structural feature involving or stabilized by the SL4b region. This additional apparent unfolding transition is also visible at low salt, but is likely stabilized beyond the measurable temperature range upon addition of Mg^2+^ (**Supplemental Fig. S2A-C**). We similarly compared the isolated SL4 and SL4-4b regions under the same conditions and observed similar major apparent unfolding transitions for each RNA (**Supplemental Fig. S2D-F**). However, under the high salt with Mg^2+^ conditions, SL4-4b exhibits an additional apparent unfolding transition at lower temperature (T_m_ ∼43°C; **Fig. 5F**). These observations suggest that SL4b influences the structure of the SL1-4 region, correlating with higher OAS1 activation, in a manner that is dependent on other structural features in this RNA, prompting us to further explore the contributions of SL1-SL3 within the context of the SL1-4b region.

### SL1 and SL3 are needed for full activation of OAS1 by the SL1-4b RNA

To assess the contributions of SL1-SL3 to OAS1 activation by SL1-4b RNA, we first generated two RNAs with 5’-end truncations, SL2-4b (deletion of SL1) and SL3-4b (deletion of SL1 and SL2). Interestingly, while the longer RNA (SL2-4b) is a weak activator, SL3-4b retains a higher level of intermediate activation that is more similar to SL1-4b (**Fig. 6A**). However, neither construct fully reaches the level of OAS1 activation by SL1-4b, suggesting that SL1 may positively influence OAS1 activation while SL2 potentially has the opposite impact in the absence of SL1. We next used UV thermal melting analysis of SL1, SL1-2, and SL1-3 RNAs in an effort to assess the contribution of each stem-loop structure to the unfolding profile of SL1-4b, but no major differences were observed (**Supplemental Fig. S2G-I**). Similar unfolding profiles were also obtained for SL2-4b and SL3-4b, but notably, only the latter exhibits a weak transition at ∼40°C under high-salt with Mg^2+^ (**Supplemental Fig. S2J-L**), resembling the low-temperature transition observed for SL4-4b (**Fig. 5F**). Thus, as before, the presence of SL4b and an additional apparent unfolding transition in the melting profile are correlated with increased capacity of the corresponding RNA to activate OAS1.

**Fig. 6.**
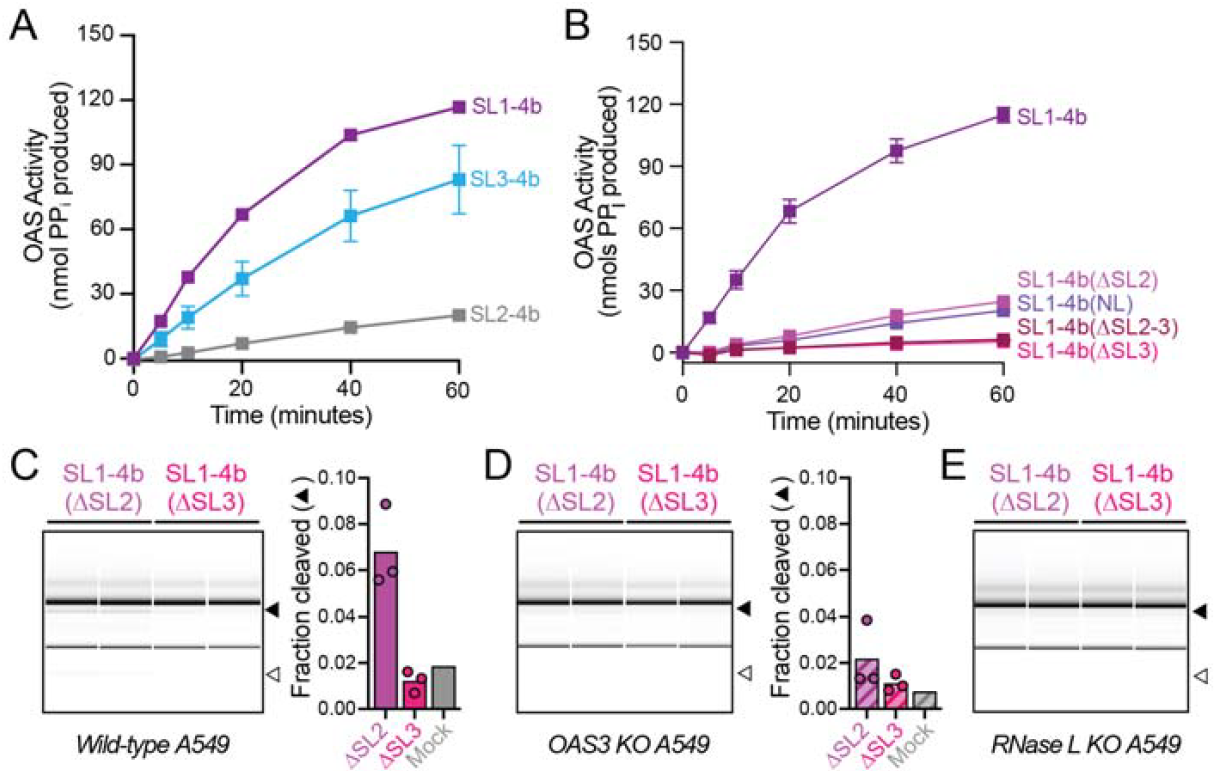
Internal SL structures (SL2 and SL3) more subtly modulate OAS1 activation by SL1-4b. *In vitro* OAS1 activation assay with **(A)** 5’-end truncations of the SL1-4b region: SL2-4b and SL3-4b, and **(B)** internal deletion variants SL1-4b(ΔSL2), SL1-4b(ΔSL3), SL1-4b(ΔSL2-3) and SL1-4b(NL). Bioanalyzer analysis of 28S and 18S integrity in **(C)** wild-type, **(D)** OAS3 KO, and **(E)** RNase L KO A549 cells. Cleavage products of 28S and 18S are shown by the black and white arrows, with cleavage of 28S being quantified for three independent transfections in wild-type and OAS3 KO cells. Quantification shows that transfection with SL1-4b(ΔSL2) leads to more cleavage of 28S by the OAS/RNase L pathway than SL1-4b(ΔSL3).

To further dissect the contributions of each internal SL structure and the linking regions between them, we generated three additional RNA constructs: SL1-4b(ΔSL2), SL1-4b(ΔSL3) and SL1-4b(NL) which lack SL2, SL3, and all single-stranded sequences between each SL (“no linker”; NL), respectively. Correct folding of each remaining SL structure within these RNAs was confirmed using SHAPE-MaP, as before (**Supplemental Fig. S3**). Again, each new RNA construct exhibited lower OAS1 activation in the *in vitro* assay than for SL1-4b, but those containing SL3, i.e., SL1-4b(ΔSL2) and SL1-4b(NL), were slightly better activators than SL1-4b(ΔSL3) which lacks this SL (**Fig. 6B**). An additional RNA construct with both SL2 and SL3 was generated (SL1-4b(ΔSL2-3), and the activity of this RNA found to match that of the RNA lacking only SL3 (**Fig. 6B**)

Finally, SL1-4b(ΔSL2) and SL1-4b(ΔSL3) were transfected into IFN-β1α-stimulated wild-type, OAS3-KO and RNase L-KO cells. In wild-type and, to a lesser extent, OAS3-KO cells, SL1-4b(ΔSL2) promotes some cleavage of 28S while SL1-4b(ΔSL3) does not appreciably activate the RNase L pathway compared to mock transfected cells (**Fig. 6C, D**). As expected, minimal rRNA cleavage was observed for either transfected RNA in RNase L KO cells (**Fig. 6E**).

### SHAPE-MaP identification of OAS1-induced changes in SL1-4b

To gain further insight into the interaction of OAS1 and the SL1-4b region and the elements within it necessary for activation, SHAPE-MaP experiments were performed for both SL1-4 and SL1-4b RNAs in the presence of equimolar (100 nM) or 10-fold excess (1 μM) OAS1. SHAPE reactivities were determined as before and values for each free RNA subtracted from those at each protein concentration to allow differences in reactivity in the presence and absence of OAS1 to be mapped onto each RNA secondary structure (**Fig. 7**). The two RNA-protein ratios were selected for these analyses as the OAS1-RNA interaction was anticipated to be potentially weak or transient in nature (23,34,35), while also recognizing that high protein concentration might lead to non-specific interactions. As such, we focused on identifying RNA regions in the more potently activating SL1-4b RNA showing consistent changes in reactivity, ideally enhanced at the higher protein concentration, rather than the absolute differences in SHAPE reactivity.

**Fig. 7.**
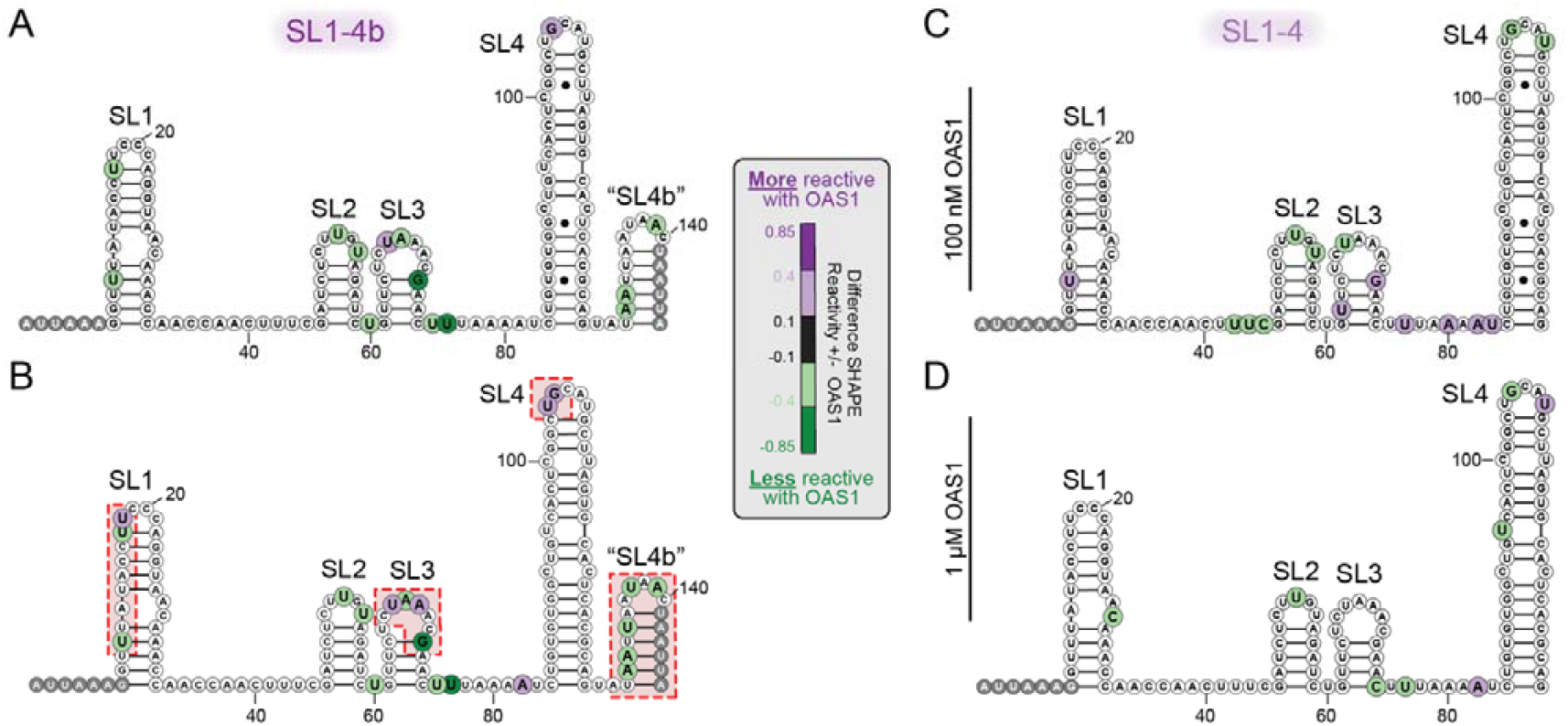
OAS1 interaction promotes changes on nucleotide dynamics in the regions identified as necessary for full OAS1 activation by SL1-4b. Average difference SHAPE reactivity (values without OAS1 subtracted from values with OAS1) from three independent modification experiments is mapped onto the RNA secondary structure for SL1-4b with **(A)** 100 nM OAS1 or **(B)** 1 μM OAS1, and for SL1-4 and **(C)** 100 nM OAS1 or **(D)** 1 μM OAS1. Changes in nucleotide dynamics are observed (red shaded regions) in SL1-4b in the regions required for full OAS1 activation, i.e. SL1, SL3, SL4 and SL4b. In contrast, SL2 exhibits similar minor differences in reactivity in both RNAs. Gray shaded nucleotides indicate primer sites for which values could not be calculated.

For SL1-4b, the most pronounced changes in SHAPE reactivity were observed at both protein concentrations at SL2 and SL4b, with the latter region showing exclusively reduced reactivity (**Fig. 7A,B**). Fewer differences were observed in SL1 (loop and stem), SL2 (loop only), and SL4 (loop only) but, where present, these were again observed consistently at both protein concentrations. In total, four regions within SL1, SL3, SL4, and 4b showed changes in reactivity at low protein concentration that became further enhanced at immediately adjacent nucleotides at high protein concentration (red shaded boxed regions in **Fig. 7B**). In contrast, for the more-weakly activating SL1-4 RNA, changes in SHAPE reactivity were less consistent at each protein concentration, with fewer nucleotides exhibiting differences at the higher OAS1 concentration. Only in SL2 were differences in reactivity similar between the two RNAs at each protein concentration. We also performed the same experiment with the full SL1-8 RNA and observed overall fewer changes in SHAPE reactivity distributed throughout the RNA secondary structure (**Supplemental Fig. S4**). As for the shorter RNAs, consistent SHAPE differences were identified in the loop of SL4 and SL4b showed exclusively reduced reactivity (albeit at fewer nucleotides). Changes in the SL1-SL3 regions all reflected increased SHAPE reactivity in the presence of OAS1, which may reflect protein-mediated disruption of additional RNA-RNA contacts in the larger RNA that are absent in the SL1-4/4b fragments.

Collectively, changes in SHAPE reactivity SL1-4 and SL1-4b RNAs support the key role of SL4b as well as the requirement of multiple other structural elements in this region in promoting optimal OAS1 activation. Finally, to gain complementary insight into the potential interactions of these RNA features with OAS1, we used AlphaFold3 (61) to model the SL1-4b-OAS1^1-346^ complex (**Supplementary Fig. S7**). All five output models place OAS1 on SL4, four position SL3 to coaxially stack with SL4, and one directly positions both SL1 and “SL4b” in contact with the C-terminal lobe of OAS1, consistent with our OAS1 activation assays and SHAPE results, as well as prior structural studies of the 5’-UTR (see below).

## Discussion

Our findings challenge the previous model that OAS1-p46 detects SL1 and/or SL2 of the SARS-CoV-2 5’-UTR to reduce disease severity (38,50). While prior studies provided an important link between OAS1 activation and the 5’-UTR, the current work demonstrates that the SL1-2 region alone is not sufficient for OAS1 activation. Instead, potent activation of OAS1 comparable to the entire 5’-SE requires SL1-4b. The ability of this 5’-SE region to activate OAS proteins *in vitro* and in cells reveals that these PRRs can detect dsRNA region(s) within architecturally complex RNAs and that structures within these regions can modulate the level of activation.

Our probing of SL1-4b shows that this full region, rather than any single stem-loop or sequence motif, is required for detection by OAS1, revealing a more structurally complex activator of OAS1 than previously identified. Additionally, analyses of the 3’-end truncations of the 5’-SE suggest that SL4 is the primary binding site for OAS1, with SL3 and SL4b also required for optimal presentation of this stem-loop for interaction with and activation of OAS1. The emerging model for OAS1 binding and activation by SL1-4b is further supported by AlphaFold3 modeling which consistently places OAS1 on SL4 (**Fig. 8** and **Supplementary Fig.S7**) and, in most models, recapitulates the coaxial stacking of SL3 and SL4, previously observed for this region in HCoV-OC43 (62). Although stem-loop regions other than SL4 are modelled by AlphaFold3 with lower confidence, at least one model is consistent with our difference SHAPE analyses, placing sites of OAS1-dependent nucleotide dynamics in SL1 and SL4b adjacent to the modeled protein (**Fig. 8A**). An indirect role for SL3 through potential coaxial stacking with SL4, as observed in HCoV-OC43 (62), may explain why SL4 is a less potent OAS1 activator in the absence of SL3. Finally, we envisage that the unstructured SL4b may play a similar, but potentially more direct, role in positioning OAS1 on SL4 (**Fig. 8B**). High-resolution structural studies, enabled by the current work’s identification of all necessary elements within the 5’-SE, will be needed to fully discern the role(s) of each SL region in OAS1 activation.

**Fig. 8.**
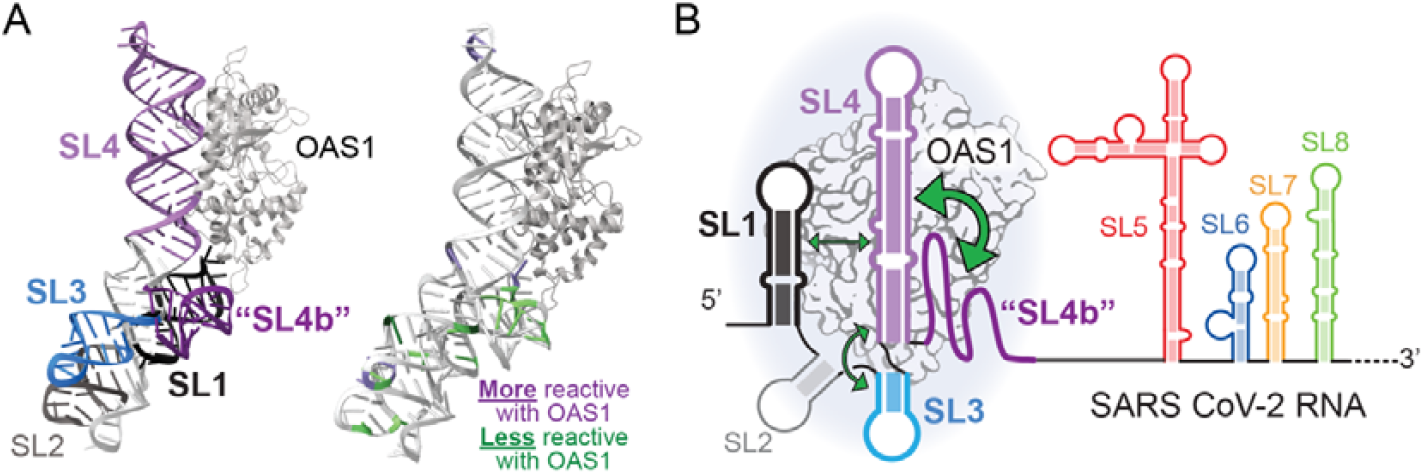
OAS1 activation by the SARS-CoV-2 5’-SL requires multiple features of the SL1-4b region. **(A)** AlphaFold3 model of OAS1 (1-346) binding SL1-4b. The left model is color-coded by stem-loops. The right model is showing difference SHAPE results from Fig.7 mapped onto RNA. OAS1 binds to SL4, which is coaxially stacking with SL3. SL1 and “SL4b” are interacting with the C-terminal region of OAS1 near the regions showing a decrease in SHAPE reactivity. **(B)** The schematic model depicts SL4 (>17 bp dsRNA) as the main OAS1 interaction site, with major (large green arrows) and supporting contributions (small green arrows) from the other RNA elements, SL4b and SL1/SL3, respectively (see main text for details).

It is also interesting that SL2 does not appear to be involved in OAS1 detection as it is the most conserved structural element in the coronaviral 5’-UTR, serving an important role in RNA replication (63-66). However, SL2 has been implicated as a determining factor for correct folding of neighboring RNA SLs as mutations lead to disruption in viral RNA architecture and therefore viral replication (65). The structural feature characterized here may be specific for betacoronaviruses that contain SL3, as this SL is not universally conserved in coronaviruses (67). The finding that SL4b lacks a defined structure and still contributes to OAS1 activation reinforces the idea that OAS1 can be regulated or modulated by non-dsRNA elements, such as the 3’-ssPy motif or the more complex structure identified within the cellular non-coding RNA nc886 (53,54). The current work thus reveals SL1-4b as a non-canonical but functional OAS1 activator and provides new insight into how RNA structural complexity, especially in viral UTRs, may be surveilled by the innate immune system.

Overall, our findings show that OAS1 recognition of viral RNA may depend more than previously appreciated on higher-order structural features that only occur in the context of larger RNA assembly. Specifically, these results raise the possibility that other OAS1 activating regions within viral RNAs may depend on multiple SLs or other structural features. Our study also focuses solely on the 5’-end of SARS-CoV-2 but leaves open questions about the role of cis-acting RNA elements that have been characterized, such as the 5’-UTR and 3’-UTR, in their interaction to circularize the genome and allow for replication (68). The identification of SL1-4b as the OAS1-activating region within the 5’-SE demonstrates that an RNA with a series of complex RNA secondary elements can activate OAS1 and sets the scene for future high-resolution structural studies to define how each element of the SL1-4b region contributes to OAS1 activation. As such, this work extends our current understanding of how the innate immune system recognizes highly structured viral RNA and could inform new approaches for targeting conserved RNA motifs that activate PRRs, potentially leading to novel therapeutic strategies in the context of coronavirus infections.

## Experimental Procedures

### RNA Synthesis

Plasmid DNA encoding nucleotides 1-3621 from strain SARS-CoV-2/human/USA/GA-PGCoE-0083E/2024 (PQ331003.1) (Genewiz) was used as template for PCR amplification of the full 5’-SE (SL1-8) region and 3’-end truncation variants (SL-7 to SL1-3) using primers listed in (**Supplemental Table S1**). PCR products were cloned into a modified pUC19 vector that also encodes a 3’-HDV ribozyme appended to the target RNA (51). Transcription templates for SL1-2 variants were cloned into the same plasmid using 5’-end phosphorylated and annealed chemically synthesized DNA strands (**Supplemental Table S1**). For SHAPE-MaP analysis, selected RNAs were additionally cloned into a plasmid encoding 5’- and 3’-end hairpin sequences and reverse transcription primer site, as described previously (69).

Plasmids were linearized by restriction enzyme digest (XbaI or XhoI) and used as templates for RNA *in vitro* transcription using T7 RNA polymerase. Reactions were incubated for 5 hours at 37 °C and subsequently quenched by addition of 250 mM EDTA followed by overnight dialysis against 1× Tris-EDTA (TE) buffer (10 mM Tris-HCl, pH 8, and 1 mM EDTA). To concentrate RNAs for purification, dialyzed RNAs were precipitated by adding 0.1 volume 3 M sodium acetate and three volumes of 100% ethanol and incubation for 1 hour or overnight at 20 °C. RNAs were spun down at 8,000 × *g* for 45 minutes at 4°C, resuspended in 1× TE and purified by urea-denaturing polyacrylamide gel electrophoresis with target RNA bands identified by UV shadowing at 254 nm, excised, and the crushed gel slices soaked in 0.3 M sodium acetate, pH 5.2 at 4 °C overnight. RNAs were recovered by ethanol precipitation and resuspension in 1× TE. RNA purity was assessed by both native and urea-denaturing gel electrophoresis (**Supplemental Fig. S1A**). SL1 RNA was obtained using commercial solid phase chemical synthesis (Integrated DNA Technologies) and used as provided after resuspension in 1x TE buffer.

### OAS1 protein expression and purification

Human OAS1 (residues 1-346) was expressed as an N-terminally tagged 6×His-SUMO fusion protein in *Escherichia coli* BL21 (DE3) as previously reported (34,60). Briefly, cells harboring the expression plasmid pE-SUMO-OAS1^1-346^ were grown in lysogeny broth at 37°C to mid-log phase (absorbance ∼0.6 at 600 nm) absorbance and expression was induced by addition of 0.5 mM isopropyl β-D-1-thiogalactopyranoside. Growth was continued overnight at 20 °C before being harvested by centrifugation and lysed by sonication in 50 mM Tris-Cl pH 8.0 buffer containing 300 mM NaCl, 20 mM imidazole, 10% glycerol and 10 mM β-mercaptoethanol. Cell lysate was cleared by centrifugation and filtration using a 0.45 μm filter before purification by Ni^2+^-affinity chromatography an ÄKTA purifier 10 FPLC. SUMO-OAS1^1-346^ fusion protein was aliquoted and stored at −80 °C in 50 mM Tris-Cl pH 7.4, 150 mM NaCl, 10% (v/v) glycerol and 2 mM dithiothreitol.

### In vitro OAS1 activation assay

OAS1 activation was assessed using an established chromogenic assay that measures pyrophosphate (PP_i_) produced during synthesis of 2’-5’ oligoadenylates (34,52,53). Before use, the 6×His-SUMO tag was removed using SUMO protease (Ulp1), expressed and purified in-house (70), and the cleaved OAS1 protein was dialyzed overnight against 25 mM Tris-HCl pH 7.4 buffer containing 50 mM NaCl, 1 mM EDTA, and 1 mM DTT. To ensure OAS1 was sufficiently pure and free of oligomeric or aggregated protein, purified protein was analyzed by SDS-PAGE before and after cleavage with SUMO protease (Ulp1) and the cleaved protein assessed by both size-exclusion chromatography on a Superdex200 10/300 column and Tycho NT.6 analysis (**Supplemental Fig. S1B-D**). RNAs were annealed in 1× TE by heating to 75 °C for 90 seconds and slowly cooling to room temperature before diluting in milli-Q water to 3 μM. Final assays contained 100 nM OAS1, 300 nM RNA, 2 mM ATP, 20 mM Tris-Cl pH 7.4, 7 mM MgCl_2_ and 1 mM DTT in a total of 150 μL and were incubated at 37 °C using a thermocycler. Control reactions included 5 μg/ mL poly(I:C) or no RNA in place of the SARS SL RNAs. Reaction aliquots (10 μL) were removed at the indicated time points and quenched by addition of 2.5 μl of 250 mM EDTA. Assays were developed by the addition of 2.5% (w/v) ammonium molybdate dissolved in 2.5 M H_2_SO_4_ (10 μL), 0.5 M β-mecaptoethanol (10 μL) and milli-Q water to a final volume of 100 μL. Absorbance at 580 nm was measured on the Synergy Neo2 (BioTek) plate reader. After subtraction of background, absorbance readings were converted to nmol PP_i_ produced using a standard curve and data plotted in GraphPad Prism 11.

### In-cell OAS/RNase L assay in A549 cells

Wild-type, OAS3 knock-out and RNase L knock-out A549 cells were constructed and validated as previously described (24). Cells were cultured in complete medium consisting of DMEM supplemented with 10% fetal bovine serum and Normocin. Prior to use, cell lines were assessed to confirm the absence of mycoplasma and were validated for RNase L or OAS3 expression by immunoblot analysis (see below). Cells were seeded in 12-well plates in complete medium and incubated at 37 °C for 7 hours to allow for attachment before treatment with 500 U/mL human IFN-β1a (PBL Assay Science, 11415-1) for 16 hours. On the day of transfection, culture medium was replaced and cells were transfected with RNA (50 nM) or poly(I:C) (0.1 µg/ml) using siLentFect and incubated at 37 °C for 6 hours. Total RNA was extracted using a RNeasy Plus miniprep kit and resolved using an Agilent 2100 Bioanalyzer system.

### Immunoblotting

Human wild-type, OAS3 knock-out, and RNase L knock-out A549 cells were either untreated or treated for 16 hours with 500 U/ml IFN-β1a (PBL Assay Science). Cells were rinsed once with phosphate-buffered saline and lysed in RIPA lysis buffer supplemented with Complete Mini protease inhibitor cocktail and clarified by centrifugation for 10 minutes (22,100 × *g* at 4 °C). Protein concentrations were determined using the Protein Assay Dye with known amounts of the provided bovine serum albumin standard. Total protein (10 μg/well) was resolved on a 4-12% Bis-Tris polyacrylamide gel and transferred to a nitrocellulose membrane at 50 V for 1 hour at room temperature. Membranes were rinsed in PBS containing 0.1% Tween-20 (PBST) and blocked in PBST containing 5% (w/v) skim milk for 1 hour at room temperature. After three washes in PBST (5 minutes), membranes were incubated in primary antibody diluted in PBST containing 5% (w/v) bovine serum albumin overnight at 4 °C. Membranes were then washed again in PBST (3 × 5 minutes) before incubating with HRP-conjugated anti-mouse (Promega #W4021, 1:2,500) or anti-rabbit (Promega #W4011, 1:2,500) secondary antibodies diluted in PBST containing 5% (w/v) skim milk for 1 hour at room temperature. Blots were washed three times in PBST (3 × 5 minutes) and then incubated in Clarity Western ECL according to the manufacturer’s recommendations and imaged using a Chemidoc MP system. The primary antibodies used in this study were: RNase L (E-9, Santa Cruz #sc-74405, mouse monoclonal, 1:1,000), OAS1 (D1W3A, Cell Signaling Technology, #14498S, rabbit monoclonal, 1:1000), OAS3 (E4X7Z, rabbit monoclonal Cell Signaling Technology, #74906S, 1:1000) and β-tubulin (9F3, Cell Signaling Technology #2128S, rabbit monoclonal, 1:1,000).

### 2′-hydroxyl acylation analyzed by primer extension and mutational profiling (SHAPE-MaP)

RNAs used for SHAPE-MaP were cloned into a structural cassette (69) and purified as described above. Purified RNAs (2 μg) were heated to 75 °C for 2 minutes and snap cooled on ice prior to refolding at 37 °C for 30 minutes in 20 mM Tris-Cl (pH 7.4) and 7 mM MgCl_2_. RNAs were treated with 2A3 (71) or DMSO at 37 °C for 5 minutes and the reaction was quenched with 1 M DTT. RNA was recovered using the NEB Monarch RNA clean-up kit and reverse transcribed with SuperScript II in a modified “MaP” buffer (25mM Tris-Cl (pH 8.0),187.5 mM KCl, 15 mM MnCl_2_, 25 mM DTT and 1.25 mM dNTPs) as previously described (72). After desalting (G-25 spin column, Cytiva), cDNA (5 µl) was PCR amplified in a 50 µl reaction using Phusion High-fidelity DNA Polymerase (New England Biolabs). For RNAs <200 nucleotides, amplicons were designed to include partial Nextera XT Illumina sequencing adapters and inline barcodes. Amplicons were purified using AMPure XP bead cleanup (Beckman Coulter) and quantified using a Qubit Flex (Invitrogen). Libraries were constructed with the Nextera XT DNA Library Preparation Kit (Illumina) and Illumina DNA/RNA UD Indexes, Tagmentation (Illumina) with 1 ng total cDNA as input (tagmentation was omitted for RNAs <200 nucleotides). The final amplified libraries were purified using AMPure XP bead cleanup. Libraries were validated by capillary electrophoresis on a TapeStation 4200 (Agilent), pooled at equimolar concentrations, and sequenced with PE100 reads on an Illumina NovaSeq 6000, targeting a depth of ∼300,000 reads per sample. Raw (RNAs >200 nucleotides) or demultiplexed (RNAs <200 nucleotides) FASTQ files were then analyzed using ShapeMapper2 (73) with default parameters and a minimum read depth of 5000. All samples passed quality control checks. Normalized nucleotide reactivities were averaged and mapped to predicted secondary structures using RNAcanvas (74).

## Abbreviations

5’-SE: 5’-end structured elements
5’-UTR: 5’-end untranslated region
dsRNA: double-stranded RNA
COVID-19: coronavirus disease 2019
HDV: hepatitis delta virus
IFN: interferon
IVT: in vitro (RNA) transcription
KO: knock out
OAS: oligoadenylate synthetase
poly(I:C): polyinosinic-polycytidylic acid
PPi: inorganic pyrophosphate
PRR: pattern recognition receptor
AMP: pathogen associated molecular pattern
SARS-CoV-2: severe acute respiratory syndrome coronavirus 2
SHAPE-MaP: 2’-hydroxyl acylation analyzed by primer extension and mutational profiling
SNP: single nucleotide polymorphism
SL: stem-loop

## Data availability

Sequencing data for SHAPE-MaP analyses are available through the NCBI Sequence Read Archive (https://www.ncbi.nlm.nih.gov/sra) under BioProject accession number PRJNA1446773. All other data are contained within the manuscript or supplemental materials.

## Supporting information

This article contains supporting information.

## Acknowledgements

We thank Drs. Anita Corbett, Christine Dunham, and Daniel Reines, as well as members of the Conn lab, for suggestions and feedback during the course of this work. We also thank Mr. Samuel Moore for technical support.

