## Supporting Information for "SARS-CoV-2 5’-UTR stem-loops activate the antiviral protein oligoadenylate synthetase 1 (OAS1)"

**This file contains**:

Supplemental Figures S1-S7
Supplemental Table S1

### SUPPLEMENTAL FIGURES

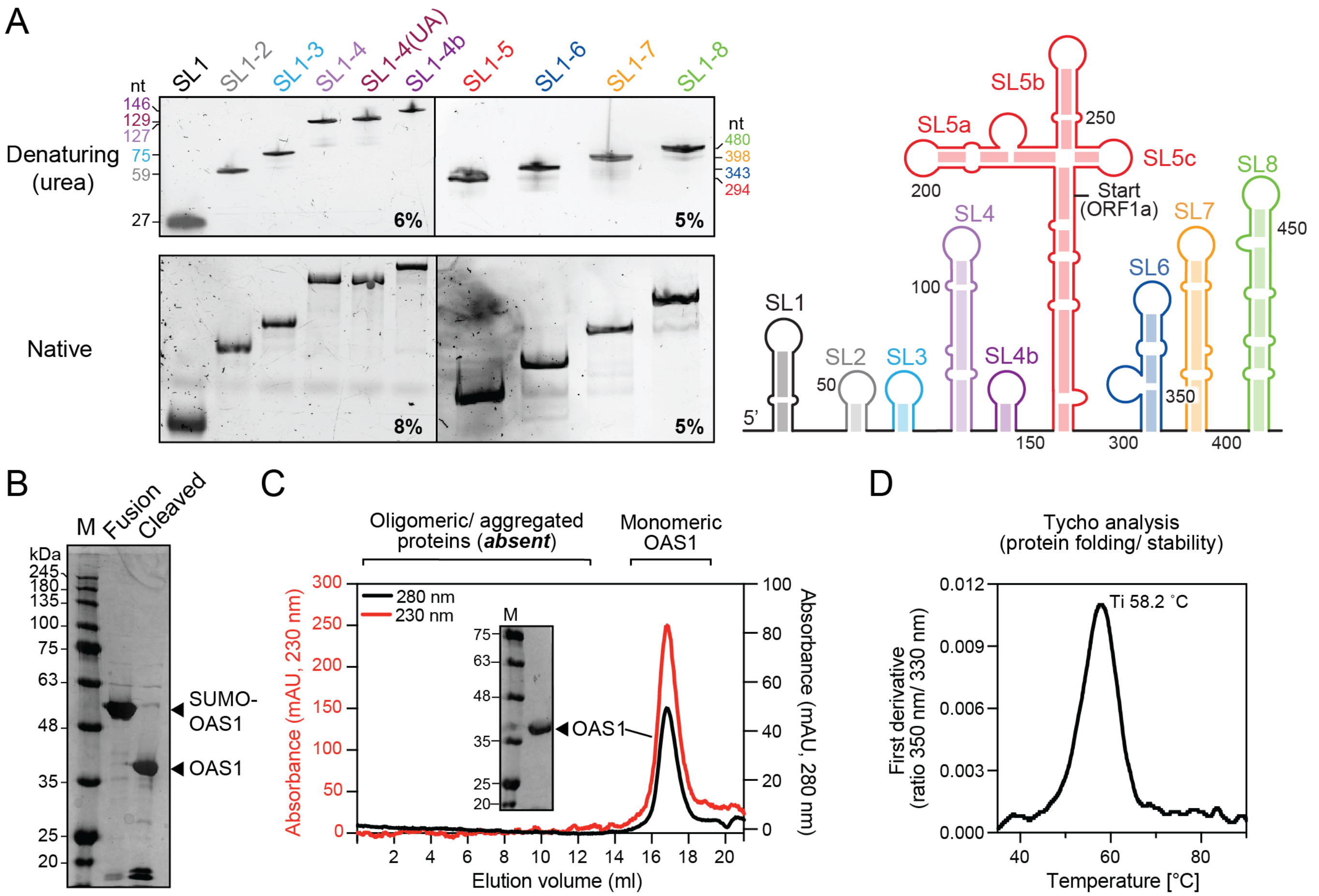


**Fig. S1. Quality control analyses of SL1-8 3'-end truncation RNAs and OAS1 protein for *in vitro* biochemical assays. (A)** *Top*: 6% polyacrylamide urea-denaturing PAGE analysis of SL1, SL1-2, SL1-3, SL1-4, SL1-4(UA) and SL1-4b RNAs with lengths in nucleotides (nt) indicated (*left)* and 5% polyacrylamide urea-denaturing PAGE of SL1-5, SL1-6, SL1-7 and SL1-8 RNAs (*right*). *Bottom*: 8% (*left*) and 5% (*right*) native-PAGE of analyses of the same sets of RNAs. **(B)** SDS-PAGE analysis of $Ni^{2+}$-affinity purified SUMO-OAS1 fusion protein before (Fusion) and after (Cleaved) treatment with SUMO protease (Ulp1). **(C)** Size exclusion chromatography analysis of OAS1 showing a single major symmetrical peak corresponding to monomeric OAS1 and the absence of oligomeric or aggregated protein. Inset shows SDS-PAGE analysis of the pooled peak fractions. **(D)** OAS1 protein folding and stability analysis using a Tycho NT.6 shown as the first derivative of the intrinsic fluorescence measurements; the calculated inflection temperature (Ti) is indicated.

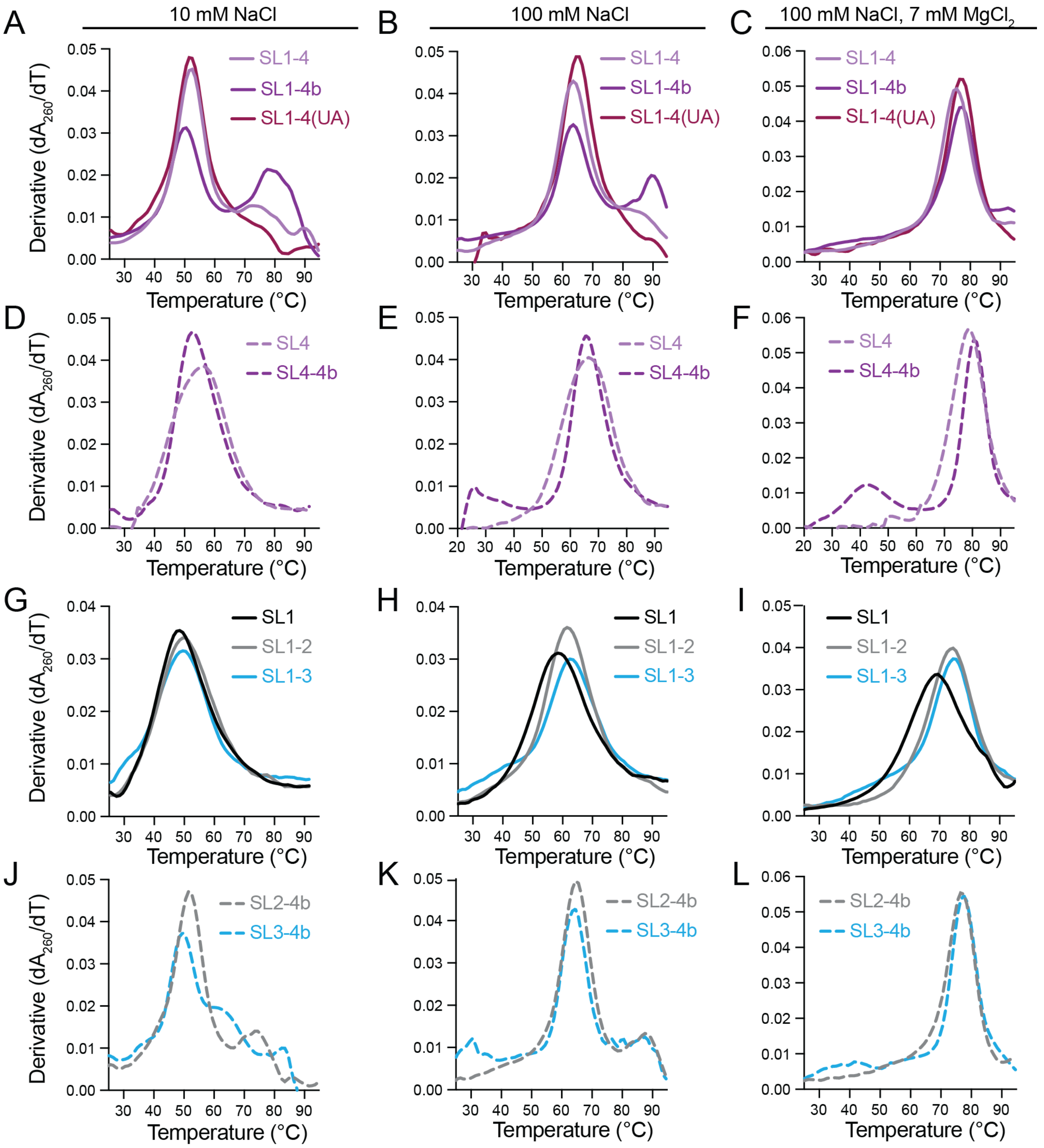


**Fig. S2. RNA UV thermal melting analyses.** Unfolding profiles (first derivative of the UV absorbance curve) are shown for SL1-4, SL1-4b and SL1-4(UA) in solution containing **(A)** 10 mM NaCl, **(B)** 100 mM NaCl, or **(C)** 100 mM NaCl and 7 mM $MgCl_2$. **(D)**-**(F)** As panels A-C but for the isolated SL4 and SL4-4b hairpin RNAs. **(G)**-**(I)** As panels A-C but for SL1, SL1-2, and SL1-3 RNAs. **(J)**-**(L)** As panels A-C but for SL2-4b and SL3-4b RNAs. The plots shown in panels B and F are the same as those in **Fig. 5E** and **5F**, respectively.

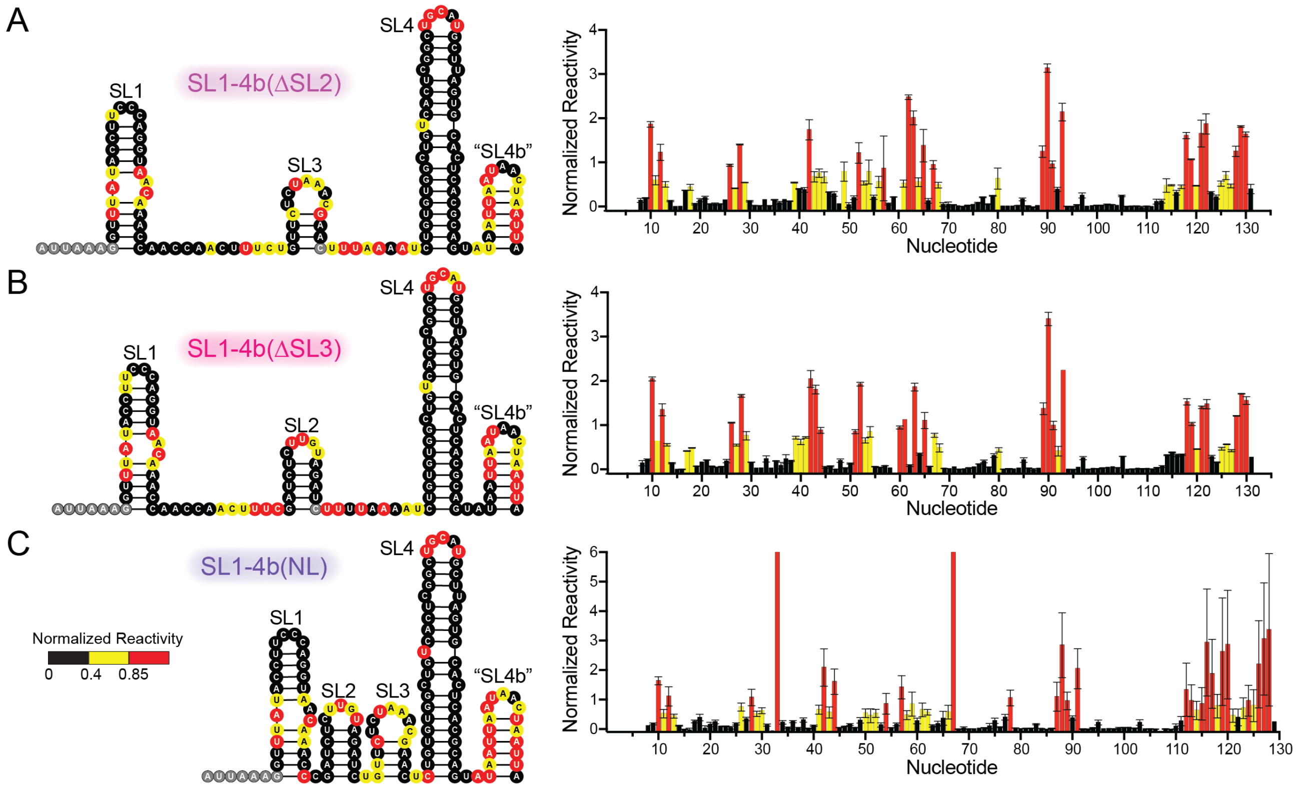


**Fig. S3. Secondary structure of SL1-4b validated by SHAPE-MaP.** Average normalized SHAPE (2A3) reactivity at 5 minutes, for two independent modification experiments mapped on the predicted secondary structure of **(A)** SL1-4b(ΔSL2), **(B)** SL1-4b(ΔSL3), and **(C)** SL1-4b(NL) RNAs. Gray shaded nucleotides indicate primer sites for which values could not be calculated. Normalized reactivity for each nucleotide is plotted on the right. SHAPE reactivity validates the structures of SL1, SL2, SL3, and SL4, however SL4b is highly modified and does not appear to form a stable secondary structure in any RNA construct.

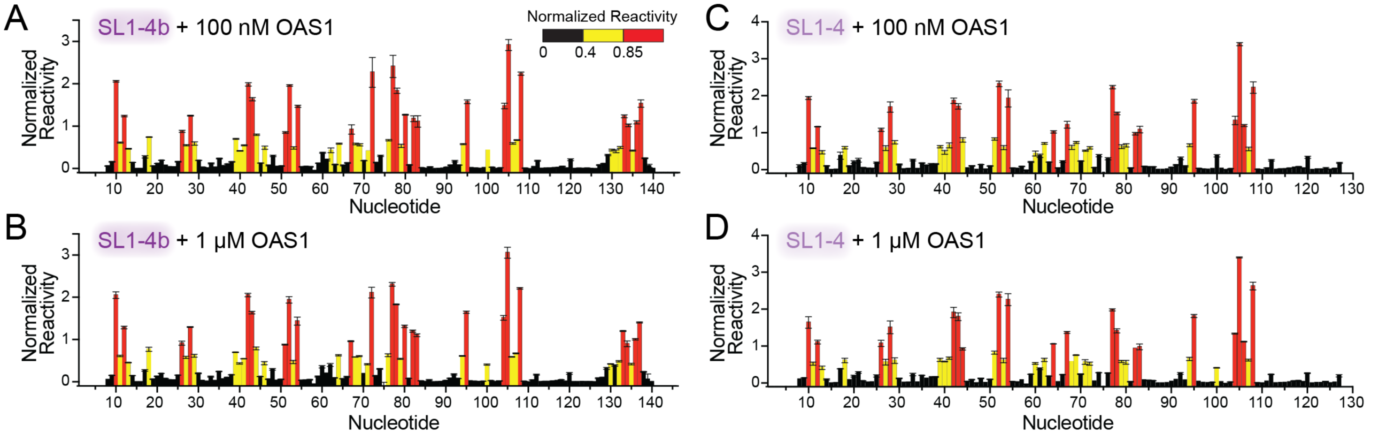


**Fig. S4. SHAPE-MaP reactivity in the presence of OAS1.** Average normalized SHAPE (2A3) reactivity at 5 minutes, for three independent modification experiments plotted for SL1-4b with **(A)** 100 nM OAS1 (1:1 protein-to-RNA ratio) and **(B)** 1 μM OAS1 (10:1 protein-to-RNA ratio), and SL1-4 with **(C)** 100 nM OAS1 (1:1 protein-to-RNA ratio) and **(C)** 1 μM OAS1 (10:1 protein-to-RNA). These data were used with SHAPE reactivities in the absence of OAS1 (**Fig. 5A,B**) to generate the difference SHAPE reactivities shown in **Fig. 7**.

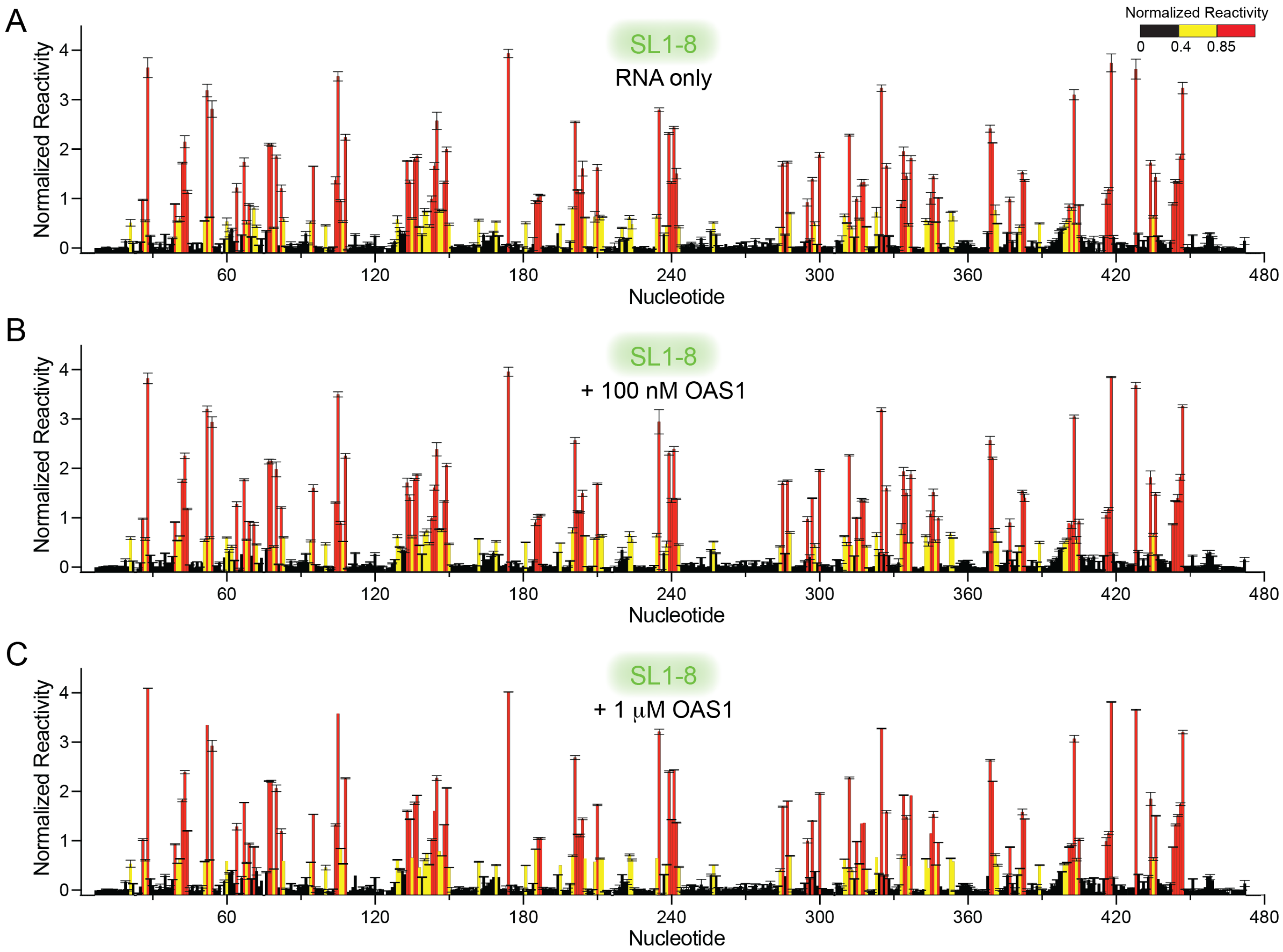


**Fig. S5. SHAPE-MaP reactivity for SL1-8 in the absence and presence of OAS1.** Average normalized SHAPE (2A3) reactivity at 5 minutes, for three independent modification experiments plotted for SL1-8 **(A)** alone or with **(B)** 100 nM (1:1 protein-to-RNA ratio) or **(C)** 1 μM OAS1 (10:1 protein-to-RNA ratio). These data were used to generate the difference SHAPE reactivities shown in **Supplemental Fig. S6**.

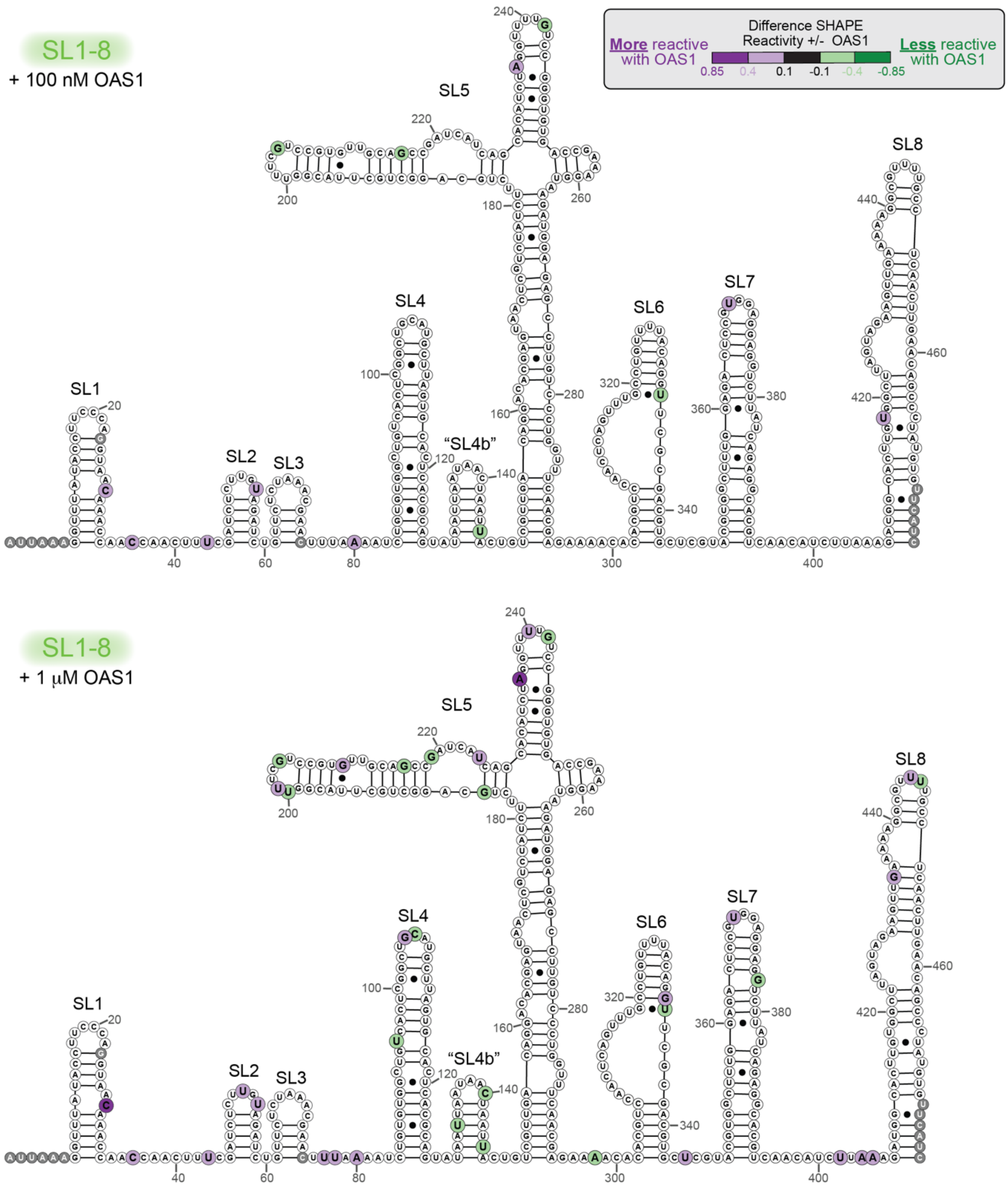


**Fig. S6. Difference in SHAPE reactivity for SL1-8 in absence and presence of OAS1.** Average difference SHAPE reactivity (values without OAS1 subtracted from values with OAS1; plotted in **Supplemental Fig. S5**) from three independent modification experiments is mapped on the SL1-8 secondary structure with 100 nM (*top*) or 1 mM OAS1 (*bottom*). Gray shaded nucleotides indicate primer sites for which values could not be calculated.

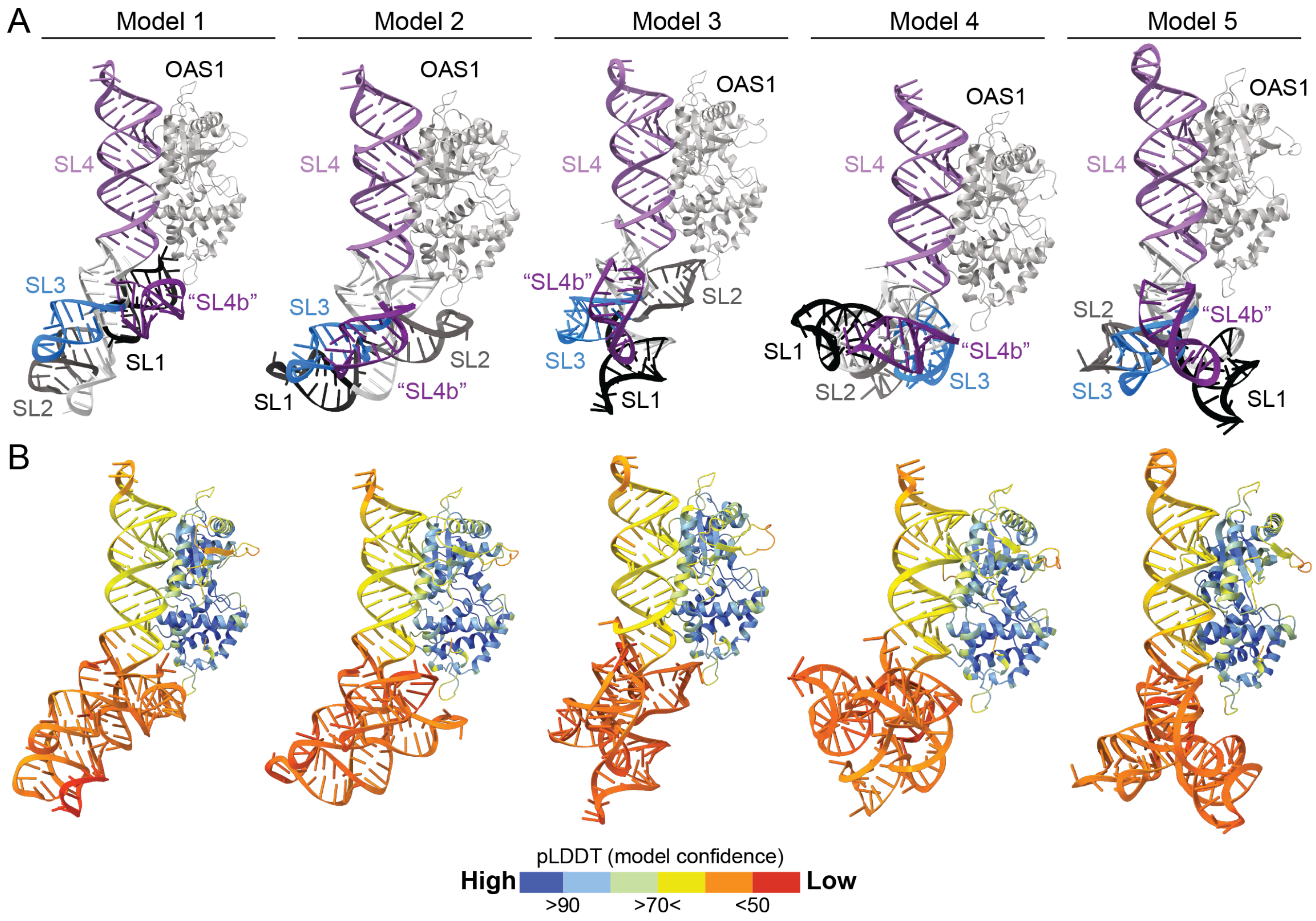


**Fig.S7. AlphaFold3 modeling of the SL1-4b-OAS1 complex. (A)** Five output models of the OAS1$^{1\text{-}346}$ interaction with SL1-4b. RNA SLs are color-coded as in the other figures. Model 1 is the same as that shown in **Fig. 8A**. **(B)** Corresponding views of each model colored according to the pLDDT model confidence values output by AlphaFold3.

### SUPPLEMENTAL TABLES

**Table S1. Sequences of DNA primers used in this study**

| Primer | Sequence 5' to 3' | Purpose |
|---|---|---|
| SL EcoRI fwd | GAATTCTAATACGACTCACTATAGG | Cloning **SL1-8** and truncated RNAs into the pUC19-HDV *in vitro* RNA transcription plasmid. |
| SL1-8 NheI rev | GCTAGCTTGATGAACACATAGGG | |
| SL1-7 NheI rev | GACTCAGCTAGCATGTTGACGTGCCTC | |
| SL1-6 NheI rev | GACTCAGCTAGCGCACGTCGCGAACCTG | |
| SL1-5 NheI rev | GACTCAGCTAGCTCGTTGAAACCAGGG | |
| SL1-4b NheI rev | GACTCAGCTAGCTAATTAGTTATTAATTATACTGCG | |
| SL1-4 NheI rev | GACTCAGCTAGCCTGCGTGAGTGCACTAAGC | |
| SL1-3 NheI rev | GACTCAGCTAGCGTTCGTTTAGAGAACAGATC | |
| SL1-2 Strand 1 | AATTCTAATACGACTCACTATAGGTTAAAGGTTTATAC CTTCCCAGGTAACAAACCAACCAACTTTCGATCTCT TGTAGATCG | Direct cloning of **SL1-2** into the pUC19-HDV *in vitro* RNA transcription plasmid. |
| SL1-2 Strand 2 | CTAGCGATCTACAAGAGATCGAAAGTTGGTTGGTTT GTTACCTGGGAAGGTATAAACCTTTAACCTATAGTGA GTCGTATTAG | |
| SL4 fwd | AATTCTAATACGACTCACTATAGGCTGTGTGGCTGT CACTCGGCTGCATGCTTAGTGCACTCACGCAGG | Deletion of SL1-3 from plasmid encoding SL1-4 to generate **SL4**. |
| SL4 rev | CTAGCCTGCGTGAGTGCACTAAGCATGCAGCCGAG TGACAGCCACACAGCCTATAGTGAGTCGTATTAG | |
| Add SL4b fwd | TATAATTAATAACTAATTACATATGCGGTTCGC | Add SL4b to constructs encoding to generate **SL2-4b**, **SL3-4b**, **SL1-4b(ΔSL2-3)**, and **SL4b**. |
| Add SL4b rev | CTGCGTGAGTGCACTAAG | |
| SL2-4 fwd | CTGACTGAATTCTAATACGACTCACTATAGATCTCTT GTAGATCTGTTCTCTAAACG | Deletion of SL1 from SL1-4 to generate SL2-4. |
| SL3-4 fwd | AATTCTAATACGACTCACTATAGTTCTCTAAACGAAC TTTAAAATCTGTGTGGCTGTCACTCGGCTGCATGCT TAGTGCACTCACGCAGG | Remove SL1-2 from SL1-4 to generate SL3-4. |
| SL2-4/3-4 rev | CTAGCCTGCGTGAGTGCACTAAGCATGCAGCCGAG TGACAGCCACACAGATTTTAAAGTTCGTTTAGAGAA CTATAGTGAGTCGTATTAG | Reverse primer used with SL2-4 fwd and SL3-4 fwd. |
| SL1+4 fwd | GAAAGTTGGTTGGTTTGTTAC | Remove SL2-3 from SL1-4 to generate SL1-4(ΔSL2-3). |
| SL1+4 rev | CTGTGTGGCTGTCAC | |
| Barcode 1 fwd | TCGTCGGCAGCGTCAGATGTGTATAAGAGACAGGG ACTCCTGCGGTTCGCCGCGGATTAAAG | Amplification primers for SHAPE-MaP: addition of barcodes and partial Illumina adaptors to cDNA.<br><br>(Used with RT SHAPE 1) |
| Barcode 1 rev | GTCTCGTGGGCTCGGAGATGTGTATAAGAGACAGA AGGAGTAGAACCGGACCGAAGCCCG | |
| Barcode 2 fwd | TCGTCGGCAGCGTCAGATGTGTATAAGAGACAGTA GGCATGGCGGTTCGCCGCGGATTAAAG | |
| Barcode 2 rev | GTCTCGTGGGCTCGGAGATGTGTATAAGAGACAGC TAAGCCTGAACCGGACCGAAGCCCG | |
| Barcode 3 fwd | TCGTCGGCAGCGTCAGATGTGTATAAGAGACAGCT CTCTACGCGGTTCGCCGCGGATTAAAG | |
| Barcode 3 rev | GTCTCGTGGGCTCGGAGATGTGTATAAGAGACAGC GTCTAATGAACCGGACCGAAGCCCG | |
| Barcode 4 fwd | TCGTCGGCAGCGTCAGATGTGTATAAGAGACAGCG AGGCTGGCGGTTCGCCGCGGATTAAAG | |
| Barcode 4 rev | GTCTCGTGGGCTCGGAGATGTGTATAAGAGACAGT CTCTCCGGAACCGGACCGAAGCCCG | |
| Barcode 5 fwd | TCGTCGGCAGCGTCAGATGTGTATAAGAGACAGAA GAGGCAGCGGTTCGCCGCGGATTAAAG | |

| | | |
|---|---|---|
| Barcode 5 rev | GTCTCGTGGGCTCGGAGATGTGTATAAGAGACAGC TCTCTATGAACCGGACCGAAGCCCG | Amplification primers for SHAPE-MaP: addition of barcodes and partial Illumina adaptors to cDNA. (Used with RT SHAPE 1) |
| Barcode 6 fwd | TCGTCGGCAGCGTCAGATGTGTATAAGAGACAGGT AGAGGAGCGGTTCGCCGCGGATTAAAG | |
| Barcode 6 rev | GTCTCGTGGGCTCGGAGATGTGTATAAGAGACAGT ATCCTCTGAACCGGACCGAAGCCCG | |
| Barcode 7 fwd | TCGTCGGCAGCGTCAGATGTGTATAAGAGACAGGC TCATGAGCGGTTCGCCGCGGATTAAAG | |
| Barcode 7 rev | GTCTCGTGGGCTCGGAGATGTGTATAAGAGACAGG TAAGGAGGAACCGGACCGAAGCCCG | |
| Barcode 8 fwd | TCGTCGGCAGCGTCAGATGTGTATAAGAGACAGAT CTCAGGGCGGTTCGCCGCGGATTAAAG | |
| Barcode 8 rev | GTCTCGTGGGCTCGGAGATGTGTATAAGAGACAGA CTGCATAGAACCGGACCGAAGCCCG | |
| Barcode 9 fwd | TCGTCGGCAGCGTCAGATGTGTATAAGAGACAGAC TCGCTAGCGGTTCGCCGCGGATTAAAG | |
| Barcode 9 rev | GTCTCGTGGGCTCGGAGATGTGTATAAGAGACAGT CGACTAGGAACCGGACCGAAGCCCG | |
| Barcode 10 fwd | TCGTCGGCAGCGTCAGATGTGTATAAGAGACAGGG AGCTACGCGGTTCGCCGCGGATTAAAG | |
| Barcode 10 rev | GTCTCGTGGGCTCGGAGATGTGTATAAGAGACAGT TCTAGCTGAACCGGACCGAAGCCCG | |
| Barcode 11 fwd | TCGTCGGCAGCGTCAGATGTGTATAAGAGACAGGC GTAGTAGCGGTTCGCCGCGGATTAAAG | |
| Barcode 11 rev | GTCTCGTGGGCTCGGAGATGTGTATAAGAGACAGC CTAGAGTGAACCGGACCGAAGCCCG | |
| Barcode alt rev 1 | GTCTCGTGGGCTCGGAGATGTGTATAAGAGACAGC TCTCTATGAACCGCATATGTAATTA | Amplification primers for SHAPE-MaP: addition of barcodes and partial Illumina adaptors to cDNA. (Used with RT SHAPE 2) |
| Barcode alt rev 2 | GTCTCGTGGGCTCGGAGATGTGTATAAGAGACAGT ATCCTCTGAACCGCATATGTAATTA | |
| Barcode alt rev 3 | GTCTCGTGGGCTCGGAGATGTGTATAAGAGACAGG TAAGGAGGAACCGCATATGTAATTA | |
| Barcode alt rev 4 | GTCTCGTGGGCTCGGAGATGTGTATAAGAGACAGA CTGCATAGAACCGCATATGTAATTA | |
| Barcode alt rev 5 | GTCTCGTGGGCTCGGAGATGTGTATAAGAGACAGA AGGAGTAGAACCGCATATGTAATTA | |
| Barcode alt rev 6 | GTCTCGTGGGCTCGGAGATGTGTATAAGAGACAGC TAAGCCTGAACCGCATATGTAATTA | |
| Barcode alt rev 7 | GTCTCGTGGGCTCGGAGATGTGTATAAGAGACAGC GTCTAATGAACCGCATATGTAATTA | |
| Barcode alt rev 8 | GTCTCGTGGGCTCGGAGATGTGTATAAGAGACAGT CTCTCCGGAACCGCATATGTAATTA | |
| Barcode alt rev 9 | GTCTCGTGGGCTCGGAGATGTGTATAAGAGACAGT TCTAGCTGAACCGCATATGTAATTA | |
| RT SHAPE 1 | GAACCGGACCGAAGCCCG | Reverse transcription primers for SHAPE-MaP |
| RT SHAPE 2 | GAACCGCATATGTAATTAG | |
| SL1-8 SHAPE fwd | GGATTAAAGGTTTATACCTTCC | Primers for SHAPE-MaP with tagmentation |
| SL1-8 SHAPE rev | TTGATGAACACATAGGGC | |
